# sfate: Schur-free, target-conditioned absorption probabilities for scalable single-cell fate analysis

**DOI:** 10.64898/2026.09.19.752930

**Authors:** Wei Zeng, Lei Shi

## Abstract

**Motivation:** Single-cell fate mapping at atlas scale is constrained by memory: in the tested CellRank 2.3.2 pip environment without the optional PETSc/SLEPc stack, the GPCCA estimator fell back—after a warning—to a dense Brandts Schur routine. Across three measured sizes, its peak memory was consistent with ≈72·n² bytes, a model that back-calculates to ≈405 GB [BACK-CALCULATED] for the 74,984-cell dataset studied here—beyond any consumer workstation.

**Results:** We present scalable-fate (sfate), which computes absorption probabilities to annotation-defined target states—where ’fate probability’ denotes absorption probability toward user-defined target states on a latent-space transition graph, conceptually related to Palantir-style diffusion-based Markov-chain fate modeling rather than a velocity-derived lineage probability—from latent-space kNN graphs via column-wise GMRES solves, without any Schur decomposition. Memory scaled linearly with cell number over four measured graph-construction tiers (R² = 0.9985; 4.03 GiB at 500,000 synthetic cells in 66.2 s); the 500k absorption solve did not converge under the default configuration, so the largest measured full-pipeline scale is the 75k real-data run (1.76 GiB). Solutions agree with float64 direct references within the prespecified 1e-5 tolerance at every tested configuration (production default L∞ = 1.74e-06; best tested configuration 3.21e-07, 31× below the threshold), and reproduce CellRank’s fate probabilities when the transition kernel and terminal representative cells are held identical. On 75k mouse 5xFAD/Cd28-cKO microglia (Ayata et al., 2025; GEO: GSE296768), the full analysis ran at 1.76 GiB peak memory, returning a six-state annotation-defined absorption landscape where the archived CellRank analysis returned three comparable arms; a controlled 2×2 comparison identifies terminal-state selection as an important contributor to the cross-tool discrepancy.

**Availability:** sfate is available at https://github.com/Ericleo-zeng/sfate (release v1.0.0, commit 5472f62; MIT license) and is archived at Zenodo (DOI: 10.5281/zenodo.22851715). The repository includes the source code, the validation suite (32 unit tests, deterministic under fixed seed), all benchmark and figure-generation scripts, the synthetic fixtures, and the records underlying every figure and table.

## Introduction

Charting how individual cells choose their fates is a central task of single-cell biology. RNA velocity first made directionality measurable from snapshot data [lamanno2018velocity, bergen2020scvelo], and a generation of computational frameworks has since turned directional signals into quantitative fate probabilities. CellRank established itself as a standard framework for this task, combining transition kernels with Markov-chain machinery—most prominently the GPCCA estimator, which identifies macrostates and terminal states and computes fate probabilities toward them [lange2022cellrank, weiler2024cellrank2]. The absorption-probability formulation it solves is shared by an established lineage of methods, including waypoint-based approaches such as Palantir [setty2019palantir], random-walk trajectory inference at atlas scale such as VIA [stassen2021via], and supervised fate-bias methods such as FateID [herman2018fateid].

What has not scaled gracefully is the data itself. Atlases of several hundred thousand to over a million cells are now routine, and practitioners regularly encounter the practical ceiling of the standard toolchain. Public issue reports provide two historical, implementation-specific reference points. In issue #540, a maintainer reported approximately 35 GiB for a 100k-cell GPCCA-plus-fate run^1^ [THIRD-PARTY HISTORICAL]; this is a third-party historical measurement rather than a benchmark from the present study. Issue #1146 concerns a different path: materialization of a dense moscot optimal-transport coupling (73,131 cells) when constructing a RealTimeKernel². We therefore do not use either issue as evidence for the 72·n^2^ Brandts relationship, which is derived only from our own measurements (Results 1).

Two constraints frame this study and, we argue, much of the single-cell community’s daily reality. First, consumer-grade hardware: a typical analysis workstation offers on the order of 30 GiB of RAM, so any method whose memory grows quadratically in cell count is not merely slow but unreachable at biologically relevant scales. Second, restricted compute environments arise for several reasons, including institutional data-governance policies for patient-derived cohorts and the limited availability of large-memory nodes. The dataset analyzed here is a mouse 5xFAD/Cd28-cKO microglia atlas and does not itself test a patient-data-governance constraint; it instead provides a realistic atlas-scale workload on locally managed hardware. Methods that are numerically validated and linear in memory on owned hardware are therefore not an optimization but an enabling condition.

We make four engineering and validation contributions. (1) We measure and delimit the dense Brandts fallback under CellRank 2.3.2 and contrast it with a PETSc/SLEPc krylov-Schur control, and thereby decouple fate-probability estimation from GPCCA’s automatic coarse-graining (Results 1). (2) We implement a Schur-free, sparse reference workflow and validate its solver against float64 direct and CellRank references on matched inputs (Results 2–3). (3) We use a controlled kernel-by-terminal-set comparison to assess where cross-workflow discrepancies enter (Results 4). (4) We report a scale-dependent solver observation—aggregate-after-solve—that motivates a factorial follow-up (Results 5). CellRank itself already supports explicit terminal-state assignment (set_terminal_states) and computes fate probabilities via sparse iterative solvers; our contribution is therefore not to introduce either explicit targets or iterative absorption solving. sfate instead makes this decomposition the primary computational architecture: when target states are supplied externally, the spectral state-discovery stage—Schur decomposition, macrostate inference, and metastability-based curation—is not required for the absorption computation, and removing it yields a pure-scipy, linear-memory, numerically certified pipeline. The remainder of the paper is organized as follows: Results 1 characterizes the memory wall; Results 2 presents sfate and its scaling behavior; Results 3 validates numerical agreement; Results 4 analyzes terminal-state curation; Results 5 reports the full 75k real-data case. Figure 0 provides a conceptual decomposition of the two pipelines.

**Figure 0.**
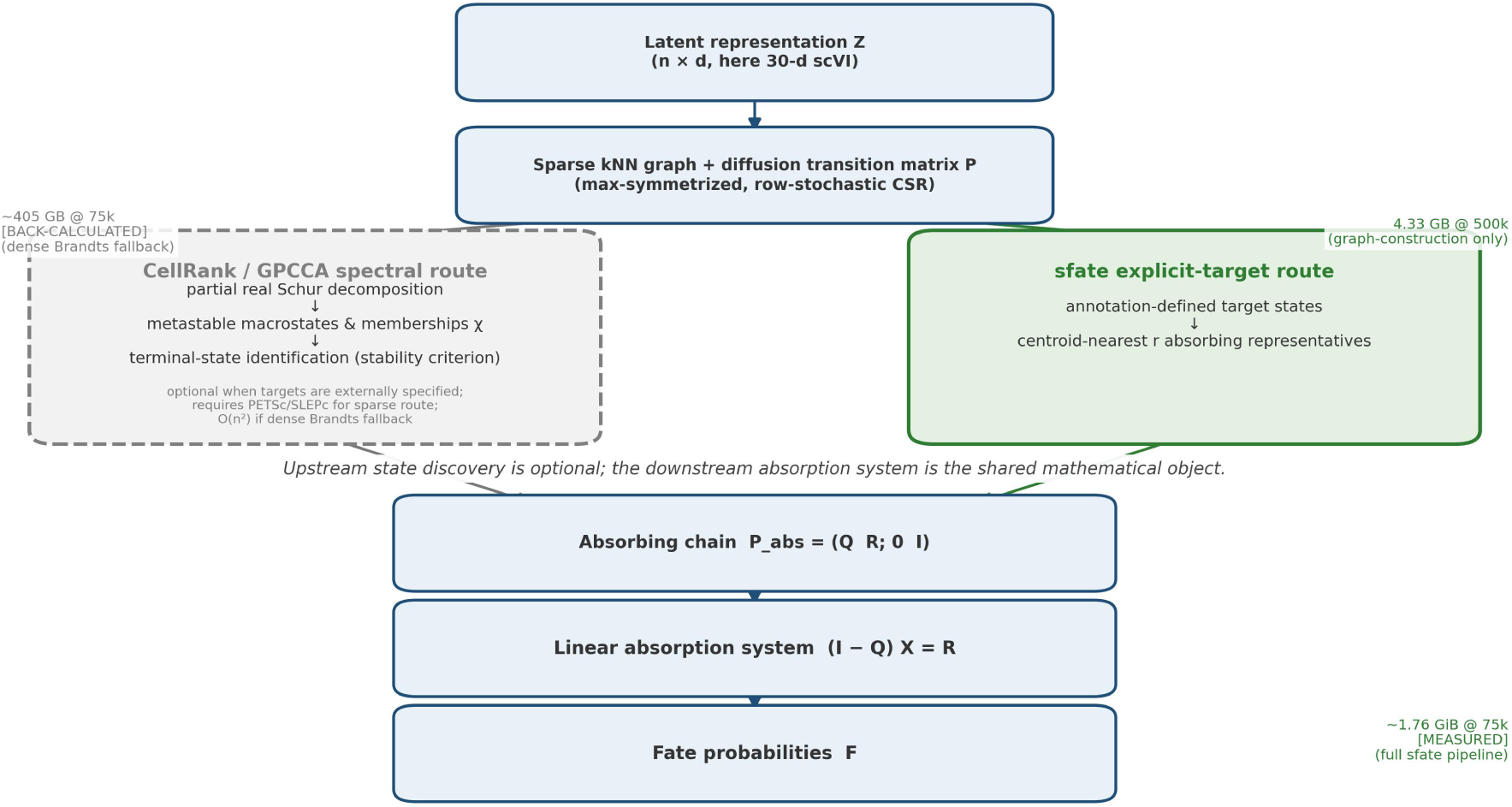
(graphical abstract). Conceptual decomposition of the two pipelines. Both routes start from the same latent representation and diffusion transition matrix. The CellRank / GPCCA spectral route (dashed) performs partial real Schur decomposition, macrostate inference, and terminal-state curation; it is optional when targets are supplied externally, and its dense Brandts fallback back-calculates to ∼405 GB at 74,984 cells [BACK-CALCULATED]. The sfate route (solid) uses annotation-defined target states and centroid-nearest absorbing representatives, then joins the same downstream absorbing-chain formulation P_abs = (Q R; 0 I), linear system (I−Q)X = R, and fate probability output F. The 500k graph-construction benchmark peaked at 4.33 GB (4.03 GiB), while the 75k full sfate pipeline peaked at ∼1.76 GiB [MEASURED].

## Results 1 — The quadratic memory wall of the default-installation Brandts fallback, not of GPCCA itself

CellRank has become a standard framework for single-cell fate mapping, and its GPCCA estimator—built on generalized Perron cluster analysis—underpins its terminal-state identification and downstream fate-probability computation [lange2022cellrank, weiler2024cellrank2]. The first computational step of this estimator is a real Schur decomposition of the n×n transition matrix. CellRank’s default Schur method is ‘krylov’, an iterative sparse path that requires the optional PETSc/SLEPc stack. In a default ‘pip install cellrank’ environment, however, PETSc/SLEPc is absent, and the estimator **automatically falls back— accompanied by a warning**—to ‘method=’brandts’’ (CellRank 2.3.2, pinned source commit 4719939, ߢestimators/mixins/decomposition/_schur.py:157–166’; pyGPCCA installation documentation describes the same full-Schur/Brandts path and its cubic scaling). For a sparse input, this fallback path densifies the transition matrix with ‘.toarray()’ and calls the LAPACK full Schur routine via the Brandts sorting algorithm [brandts2002schur, anderson1999lapack].

We measured the peak memory of this fallback path in a controlled sandbox (cellrank 2.3.2): 268 MiB at n=2,000, 1.68 GiB at n=5,000, and 4.29 GiB at n=8,000; the 2k/5k points were reproduced on the cellrank 2.1.0 installation used throughout this study to within RSS sampling precision (288.3 vs 288.2 MB; 1,800.6 vs 1,800.3 MB). The three measured points are **consistent with** an empirical n² memory model with a coefficient of ≈72 B per cell² **over the tested range** (72.1 / 72.0 / 72.0 B·n⁻² respectively)—an empirical description for this matrix class and version, not a universal law (the dense matrix itself, 8·n² bytes, is only the lower bound; the remaining factor reflects additional allocations in this execution path, whose individual contributions were not separately profiled). At n=10,000 the process was killed by the OOM killer in a 4 GB sandbox—a censored observation, not a scaling point. The model back-calculates the requirement at n=100,000 as ∼671 GiB and at n=500,000 as ∼16.4 TiB [EXTRAPOLATED from the three measured points].

For a real-world reference point, we considered the 74,984-cell microglia dataset analyzed in this study. Substituting n=74,984 into the calibrated model gives 72×74,984² B ≈ **405 GB [BACK-CALCULATED]** (≈377 GiB)—≈13.5× the ∼30 GiB physical RAM of a typical single-cell analysis workstation. In other words, under a default installation, the dataset that motivates this paper is physically infeasible on consumer-grade hardware, and cloud workarounds are not always available when data cannot leave institutional infrastructure.

This is not a hypothetical edge case. The original CellRank analysis of this dataset (the "step12" run of our Alzheimer’s disease pipeline) only succeeded after **three manual patches**: (i) restricting the VelocityKernel to a 1,673-gene subset (down from 17,714 genes; the four dense n×genes float64 arrays would otherwise have demanded ∼42 GB, reduced to ∼4 GB by the patch); (ii) forcing ‘use_petsc=Falsè with an ILU-preconditioned scipy GMRES for fate probabilities; and (iii) a manual ‘conda install petsc4py slepc4py’ on 2026-09-08 02:56, after the OOM-era attempts (backup of the OOM-version script timestamped 02:01) and 25 minutes before the first successful run at 03:21. The available provenance—‘conda-meta/history’, script timestamps, and PETSc signal-handler noise in the archived run logs—is **consistent with** the successful step12 run having used the krylov-Schur path. Because the run was not re-executed under a frozen environment, we treat this as a **provenance-based inference** rather than a direct benchmark: at 75k, the brandts path is physically impossible on a 30 GB machine, but we do not claim command-level certainty. The recurrence of the terminal-state curation artifacts (Results 4) in that archived output therefore argues against, but does not by itself exclude, an implementation-specific contribution from the Schur route; since the pattern also occurs under our measured krylov-Schur controls, it is best read as a property of the coarse-graining layer rather than of the dense fallback.

To delineate the scope of the wall precisely, we benchmarked the krylov-Schur path directly on synthetic branching topologies with PETSc/SLEPc available (petsc4py 3.25.5 / slepc4py 3.25.1, cellrank 2.1.0, single process, BLAS threads = 1) [hernandez2005slepc, stewart2002krylovschur]. At n=100,000 the full GPCCA pipeline (neighbors → ConnectivityKernel → krylov-Schur → macrostates → terminal states → fate probabilities) completed in 2 min 3 s with a whole-process peak RSS of **1.53 GiB**; the Schur step itself took 5.22 s with a +247.7 MB stage increment. At n=500,000 the same pipeline completed in 7 min 31 s with a peak RSS of **3.67 GiB** (Schur step 25.91 s, +1.24 GB increment). Both runs exited cleanly with zero swap usage. The quadratic wall is therefore **a property of the default installation’s Brandts fallback, not a mathematical upper bound of GPCCA**: with the heavy PETSc/SLEPc stack correctly installed, the wall dissolves at least to the half-million-cell scale. Figure 1 summarizes these three regimes on a common scale: the empirically calibrated n² brandts model (with the 405 GB back-calculation at 75k), the measured krylov-Schur points (1.53 GiB @ 100k, 3.67 GiB @ 500k), and the linear-memory behavior of our sfate implementation (4 tiers, 0.71–4.03 GiB), to which we turn in Results 2.

**Figure 1.**
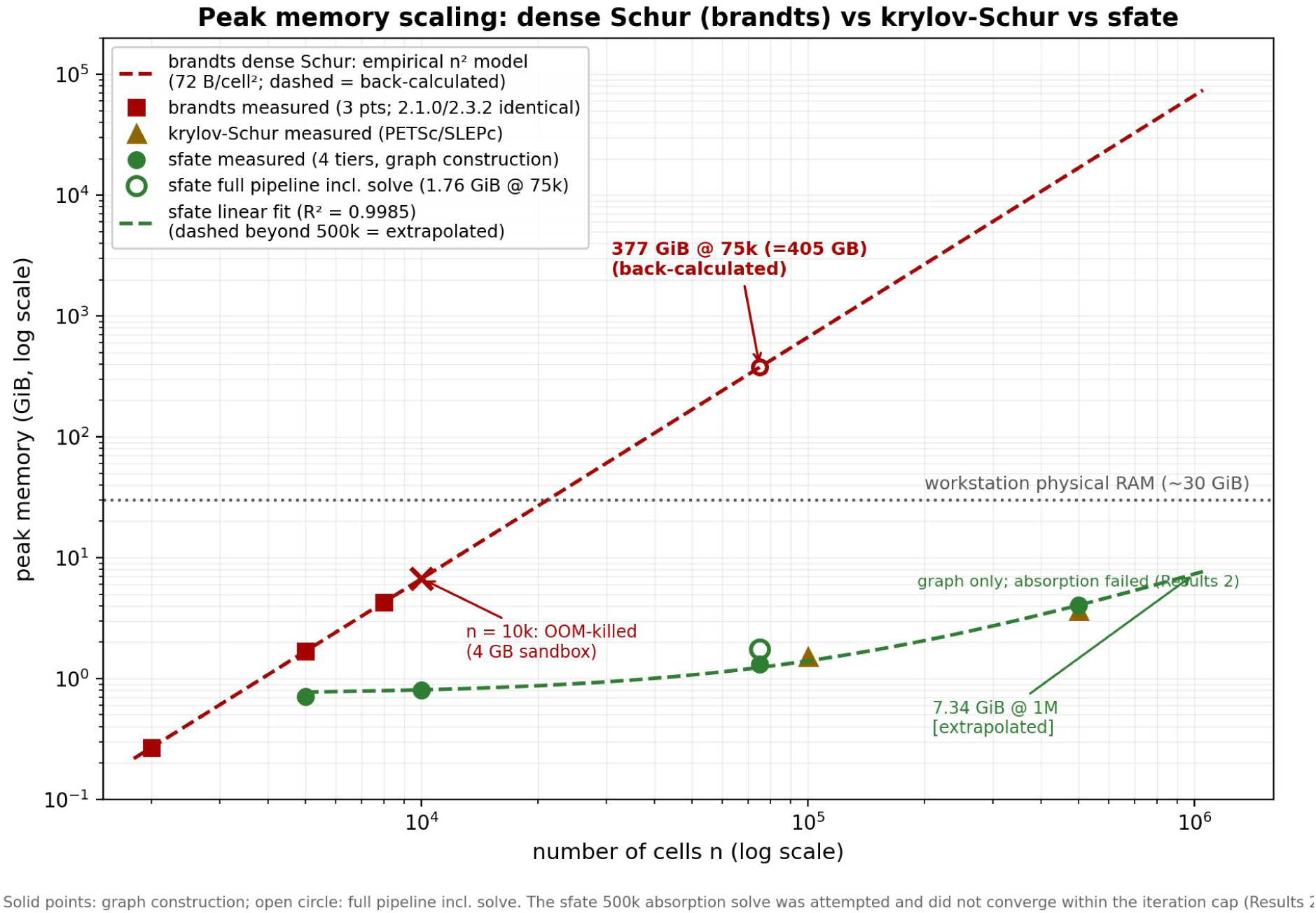
Peak memory scaling: dense Schur (Brandts fallback, no PETSc/SLEPc) vs krylov-Schur (PETSc/SLEPc) vs sfate (log-log). Marker classes distinguish [MEASURED] points, the [BACK-CALCULATED] 75k Brandts value (405 GB), and [EXTRAPOLATED] values (open/hatched symbols). The sfate linear fit has R² = 0.9985 over the four measured graph-construction tiers. Units: GB = 10⁹ bytes; GiB = 2³⁰ bytes. The sfate 500k point is the graph-construction benchmark; a full-pipeline 500k run under default column-wise GMRES did not converge (49,904 iterations, true residual 1.09e-5 at rtol = 1e-6), so the 500k absorption solve is reported as a negative result.

These three regimes summarize the memory landscape of single-cell fate computation (Figure 1, Table 1). **Regime I—default CellRank**: the Brandts fallback, whose empirically calibrated n² model places a real 75k analysis at ≈405 GB [BACK-CALCULATED]—out of reach on consumer hardware. **Regime II— properly provisioned CellRank**: the PETSc/SLEPc krylov-Schur path, which we measured to complete the full GPCCA pipeline at 100k and 500k cells within single-figure GiB, at the price of a heavy dependency stack. **Regime III—sfate**: a Schur-free absorption solver with linear memory, light dependencies, an explicit numerical validation protocol, and annotation-controlled target semantics. The contribution is therefore not to make GPCCA scalable where it is mathematically impossible, but to provide a lightweight fate-computation route that remains scalable without the optional eigensolver stack and without GPCCA’s automatic terminal-state curation.

**Table 1.** Peak memory and wall time across methods and scales

| Method/route | $n$ | Peak memory | Wall time | Evidence class and notes |
| --- | --- | --- | --- | --- |
| CellRank brandts fallback (default install) | 2,000 | 268 MiB | 7.81 s (Schur) | [MEASURED] $n^2$ calibration point; cellrank 2.3.2 survey sandbox; 2.1.0 reproduction 288.3 MB |
| CellRank brandts fallback (default install) | 5,000 | 1.68 GiB | 98.73 s (Schur) | [MEASURED] calibration point; 2.1.0 reproduction 1,800.6 MB; dense matrix is $8 \cdot n^2$ lower bound |
| CellRank brandts fallback (default install) | 8,000 | 4.29 GiB | 167.8 s (Schur) | [MEASURED] third point (E3, 2026-09-10, cellrank 2.1.0); coefficient $72.0 \text{ B/n}^2$ |
| CellRank brandts fallback (default install) | 10,000 | — | — | [MEASURED] censored: OOM-killed in 4 GB sandbox |
| CellRank brandts fallback | 74,984 | $\approx 377 \text{ GiB}$ (405 GB) | — | [BACK-CALCULATED] $72 \times 74,984^2 \text{ B}$; $\approx 13.5 \times 30 \text{ GiB}$ workstation |
| CellRank brandts fallback | 100,000 | $\approx 671 \text{ GiB}$ (720 GB) | — | [EXTRAPOLATED] from empirical $n^2$ model |
| CellRank brandts fallback | 500,000 | $\approx 16.4 \text{ TiB}$ (18 TB) | — | [EXTRAPOLATED] from empirical $n^2$ model |
| CellRank krylov-Schur (PETSc/SLEPc) | 100,000 | 1.53 GiB | 2:02.98 (full); Schur 5.22 s | [MEASURED] time -v authority; Swaps 0; cellrank 2.1.0 |
| CellRank krylov-Schur (PETSc/SLEPc) | 500,000 | 3.67 GiB | 7:30.64 (full); Schur 25.91 s | [MEASURED] time -v authority; Swaps 0 |
| CellRank GPCCA+fate (community report) | 100,000 | $\approx 35 \text{ GiB}$ | $\approx 2 \text{ min}$ | [THIRD-PARTY HISTORICAL] issue #540 (3 terminal states $\times 30$ representatives); not measured here |
| **sfate** (graph construction) | 5,000 (real fixture) | 0.71 GiB | 1.9 s | [MEASURED] RSS sampling peak; single process BLAS=1 |
| **sfate** (graph construction) | 10,000 (synthetic) | 0.80 GiB | 3.9 s | [MEASURED] RSS sampling peak; single process BLAS=1 |
| **sfate** (graph construction) | 74,984 (real) | 1.32 GiB | 18.4 s | [MEASURED] M1 benchmark $\geq 3$ reps maximum |
| **sfate** (75k full pipeline incl. solve) | 74,984 (real) | 1.76 GiB | graph 65.0 s + solve 29.0 s | [MEASURED] time -v approximate authority; six annotation-defined targets |
| **sfate** (graph construction) | 500,000 (synthetic) | 4.03 GiB | 66.2 s | [MEASURED] process authority 4.05 GiB/7:50; graph-construction benchmark |
| **sfate** (500k full pipeline, attempted) | 500,000 (synthetic) | 4.03 GiB | graph 66.2 s; solve not converged | [MEASURED] graph peak; column-wise GMRES hit maxiter 49,904 at column 0 (true residual $1.09\text{e-}5$); no measured solve time |
| **sfate** (extrapolation) | 1,000,000 | $\approx 7.34 \text{ GiB}$ | — | [EXTRAPOLATED] linear fit over graph-construction tiers; $\times 1.25$ safety factor = 8.96 GiB; unvalidated for absorption stage |

Version note: the empirical n² relationship was fitted on the cellrank 2.3.2 survey baseline (three measured points, reproduced on 2.1.0 to within sampling precision; third point at n=8,000 in output/brandts_8k.md); the krylov-Schur controls and all B/C-layer analyses used cellrank 2.1.0. Version-specific differences between 2.1.0 and 2.3.2 are enumerated in Methods and Supplementary Table S1.

## Results 2 — sfate: a Schur-free, linear-memory architecture for target-conditioned absorption

sfate replaces the GPCCA-centered object stack with a five-stage pipeline designed around one invariant: no object larger than O(n·k) is ever materialized. Structurally, for fixed neighbor count k, latent dimension d, and representative count r, the pipeline’s persistent structures are O(n·d) + O(n·k) + O(n·r)— never O(n²); the linear fit reported below is the empirical counterpart of this structural property, not its substitute. The input is a low-dimensional latent representation (in this study, a 30-dimensional scVI latent, (74,984 × 30) float32) [lopez2018scvi], from which an approximate kNN graph is built with NN-Descent (pynndescent 0.6.0 [dong2011nndescent], k=30, random projection trees plus graph diversification; full parameterization in Methods). The graph is converted to a row-stochastic transition matrix in a single CSR float32 copy (nnz ≈ 1.6–1.7·n·k after max-symmetrization), terminal states are supplied directly from cell-type annotations as small sets of representative absorbing cells (30 per state), and fate probabilities are obtained by solving the absorption system (I−Q)X = R column-wise with unpreconditioned GMRES [saad1986gmres]—no Schur decomposition, no macrostate identification, no PETSc/SLEPc. Densification is excluded mechanically rather than by convention: the test suite monkeypatches ‘toarray’/‘todensè to raise, and a CI grep forbids both calls in ‘src/’. At the I/O level, for this processed h5ad and the access path used in our pipeline, opening with anndata 0.12.16 in backed mode increased RSS by +6.9 GB on open; we therefore read latent and metadata directly with h5py in read-only mode. This is an implementation-specific measurement for this file, version, and access path, not a general characterization of AnnData backed mode [virshup2024anndata]; a minimal reproducer and the accessed keys are provided in the repository.

Empirically, memory scales linearly with n over the four measured tiers, spanning two orders of magnitude (single process, BLAS threads = 1, ≥3 repetitions, RSS sampling plus ‘/usr/bin/time -v’ as the process-level authority): 0.71 GiB @ 5k (real fixture), 0.80 GiB @ 10k (synthetic), 1.32 GiB @ 75k (real, full graph construction 18.4 s), and **4.03 GiB @ 500k** (synthetic, 66.2 s—graph-construction benchmark; the 500k full pipeline was attempted but the absorption solve did not converge under the default configuration, see Discussion). For the 500k full-pipeline attempt, graph construction completed at 4.03 GiB peak RSS; column-wise GMRES for the first absorbing cell reached 49,904 iterations without reaching rtol = 1e-6 (true residual 1.09e-5), so a measured end-to-end solve time at 500k is not available and the 1M extrapolation applies only to graph construction and I/O. The whole-benchmark process peak over the measured tiers was 4.05 GiB in 7 min 50 s (time -v, zero swap). A four-point linear fit over the graph-construction tiers gives **mem(n) = 740 + 6.60 MiB per 1,000 cells, R² = 0.9985**—a goodness-of-fit description over the measured range, not a proof of linearity; **no superlinear trend was resolved across the four measured tiers, although four points do not exclude such behavior outside this range**. Consistent with linearity, the per-cell footprint M(n)/n decreases monotonically from 142 to 8.1 MiB per 1,000 cells across the four tiers, converging toward the asymptotic slope. The fit extrapolates to 7.34 GiB at 1M cells (8.96 GiB with a 1.25 safety factor) [EXTRAPOLATED]; because the 500k solve failed to converge, this extrapolation is unvalidated for the absorption stage and should be treated as an upper bound for the graph-construction/I/O component only. The graph itself is cheap: the **theoretical** CSR footprint accounts for only 0.34 MiB per 1,000 cells (k=30, float32, nnz≈1.5·n·k); the **measured** slope (6.60 MiB per 1,000 cells) is ∼19.7× larger, a gap **consistent with** temporary allocations from NN-Descent index construction, kNN query buffers, and int64 key arrays used during symmetrization. Real-data hub structure (median row degree 40, maximum 389 at k=30) neither breaks the linear fit nor, as Results 3 shows, the solver.

The kNN backend was selected by measurement, not by default: scikit-learn’s exact trees win at 5k (0.18 s vs 1.56 s) and 10k (0.43 s vs 1.80 s), but at 75k pynndescent overtakes it decisively (6.4 s vs 14.9 s). Unlike the exact backend, NN-Descent completed the 75k and 500k workloads with a fixed seed (20260909) and an empirical approximation-quality behavior documented in the original algorithm paper [dong2011nndescent]; because it is approximate, we treat the 500k completion as a scalability result, not an accuracy guarantee. Neighbor recall against exact search on the tractable tiers and seed-sensitivity analysis are reported here: on the 5k real fixture, pynndescent recall was 0.9803 ± 0.0001 across seeds 20260909/20260910/20260911; on the 10k synthetic fixture, recall was 0.9945 ± 0.0000. Seed sensitivity was therefore negligible, but 500k accuracy was not evaluated because exact search is intractable at that scale. Varying k on the 5k fixture (r = 30 representatives per class) shows the fate landscape is rank-stable: Spearman correlation to k = 30 is 0.884 / 1.000 / 0.949 / 0.895 for k = 15/30/50/100 and MAE ≤ 1.6 × 10⁻². Graph-construction time is 1.1 / 1.5 / 2.6 s for k = 30/50/100 (the k = 15 run includes one-time index-build overhead and is not protocol-grade; it is recorded in the experiment log). pynndescent is therefore the default, with sklearn retained as a cross-check branch.

Because the landscape is computed on a latent representation, we tested its dependence on the embedding itself on the 5k fixture: replacing the 30-dimensional scVI latent with a 30-dimensional PCA embedding of the same cells (top 2,000 highly variable genes; fixed seed) yields median per-state Spearman 0.165 (range −0.37 to 0.64) and mean MAE 0.116 against the scVI reference under an identical protocol (k = 30, r = 30). The six-state absorption landscape is therefore not quantitatively preserved across these latent representations, while remaining conditional on the chosen embedding.

We are explicit about what sfate does **not** claim. At the largest measured scale, sfate’s 4.03 GiB @ 500k is **comparable to—not better than—** the 3.67 GiB measured for CellRank’s krylov-Schur path at the same n (a ∼10% difference, not an advantage; Results 1); sfate is not a speedup. Its advantages are four orthogonal properties: (i) **no cliff by default**—the dense-Schur fallback cannot be triggered because the Schur step does not exist, on any installation; (ii) **light dependencies**—pure scipy, pip-installable, no PETSc/SLEPc/MPI toolchain; (iii) an **explicit validation protocol**—numerical agreement with float64 direct and CellRank reference solutions within a prespecified tolerance (Results 3); and (iv) **annotation-controlled target semantics**—absorption probabilities for every annotation-defined target state without an automatic curation step whose behavior we dissect in Results 4. The full scaling comparison across the three regimes (Brandts back-calculation, krylov-Schur, sfate) is summarized in Figure 1 and Table 1.

## Results 3 — Numerical validation: agreement with float64 direct and CellRank reference solutions within a prespecified tolerance

Scalability claims are only meaningful if the computed fate probabilities are numerically trustworthy. We therefore validated the sfate solver under a three-layer protocol in which each layer has an explicit reference solution and acceptance criterion. **Layer A** (unit level) compares, on identical transition matrices and identical absorbing sets, sfate’s production path (float32 GMRES [saad1986gmres], rtol = 1e-6) against a float64 direct solve (scipy ‘spsolvè) as the golden reference. Acceptance combines two criteria with distinct roles: a **strict solver-level criterion** (backward error ≤ 1e-12, indicating that the computed iterate is consistent with the exact solution of a nearby linear system within the stated backward-error bound) and a **lenient application-level tolerance** (observed L∞ ≤ 1e-5 against the float64 reference, certifying fitness for biological interpretation); their relationship—including the condition-number-scaled forward bound linking them—and their calibration history are given in Methods [higham2002accuracy]. **Layer B** (cross-validation) runs the sfate solver on CellRank’s own transition kernel and CellRank’s own terminal representative cells, and reconciles the output against CellRank’s fate probabilities—thereby isolating solver agreement from kernel and terminal-state-set differences (which Results 4 shows are the largest contrast in this design). **Layer C** (real data) compares the full 75k sfate output against the existing step12 CellRank products as a record-style control with no pass gate, since the two sides differ by construction in their terminal-state sets.

The Layer A headline result (5k real-data fixture, six annotated states as absorbing classes, 30 representative cells each, block size swept over {1, 16, 256, full}) is numerical agreement with the float64 golden reference within a prespecified tolerance: across the full 2×2 configuration grid (solve mode × preconditioner), L∞ ranges from 2.34e-07 to 1.74e-06, all passing the 1e-5 acceptance tolerance with margins of 5.7×–42.7×; the current production default (column-wise, unpreconditioned) gives **L∞ = 1.74e-06 (5.7× within tolerance)**, and the ILU-preconditioned aggregated variant gives 3.21e-07 (31.2×). Per-class GMRES converged in 26–30 iterations with a maximum true residual of 1.4e-7 and zero float64 fallbacks; the four block sizes produced **identical stored outputs**, confirming that blocking is purely a memory-bounding device for this implementation. The same-level lumping check (a1) passed as well: per-cell absorption followed by within-class summation equals pseudo-state lumping exactly (float64 direct solve, full chain)—a direct self-contained proof of the mathematical equivalence claim for aggregate-after-solve evaluation, which we exploit again in Results 5.

Layer B cross-validation closes the loop against CellRank itself: with the CellRank kernel and CellRank terminal cells held fixed, sfate reproduces CellRank’s fate probabilities with **L∞ = 1.07e-06** (residual max 1.55e-07, zero orphan cells excluded)—the same order as the Layer A internal precision. This result supports solver agreement at the tested tolerance on identical inputs; it does not establish bitwise identity, and—because the terminal-cell sets are held fixed—it does not establish equivalence between CellRank’s terminal-set construction and annotation-based top-30 representative selection.

The two solve modes of sfate—aggregated-RHS (class-summed right-hand sides) versus column-wise (per-absorbing-cell solves with post-hoc class aggregation)—reconcile at **L∞ = 4.4e-06** (p99 = 6.9e-07). Note that this output-level difference exceeds 1e-6: the solver parameter is a residual stopping tolerance (rtol = 1e-6), not a direct bound on solution error, and the two modes were stopped independently under that criterion. We therefore report the modes as numerically close on this fixture; the per-mode true residuals and condition-scaled error bounds are reported in Methods. Their mathematical equivalence follows from linearity (superposition of solutions), and their numerical divergence at scale—aggregated right-hand sides may excite slow diffusive modes [titleypeloquin2014, sharpewales2021]—is the subject of the finding in Results 5, where we treat the mechanism as a hypothesis rather than an established cause.

On the 5k fixture, the float64 direct reference required 37.7 s for a single class (fill-in explosion), whereas the production GMRES path required **0.73 s** (37.7 / 0.73 ≈ 51.6). This comparison motivates the iterative production path at the tested scale; larger-scale direct-solve behavior was not measured, and we do not claim that the iterative route is the only practical one in general. Figure 2 summarizes the validation evidence across the four scenarios; only Layers A and B carry acceptance gates, and Layer C is record-style.

**Figure 2.**
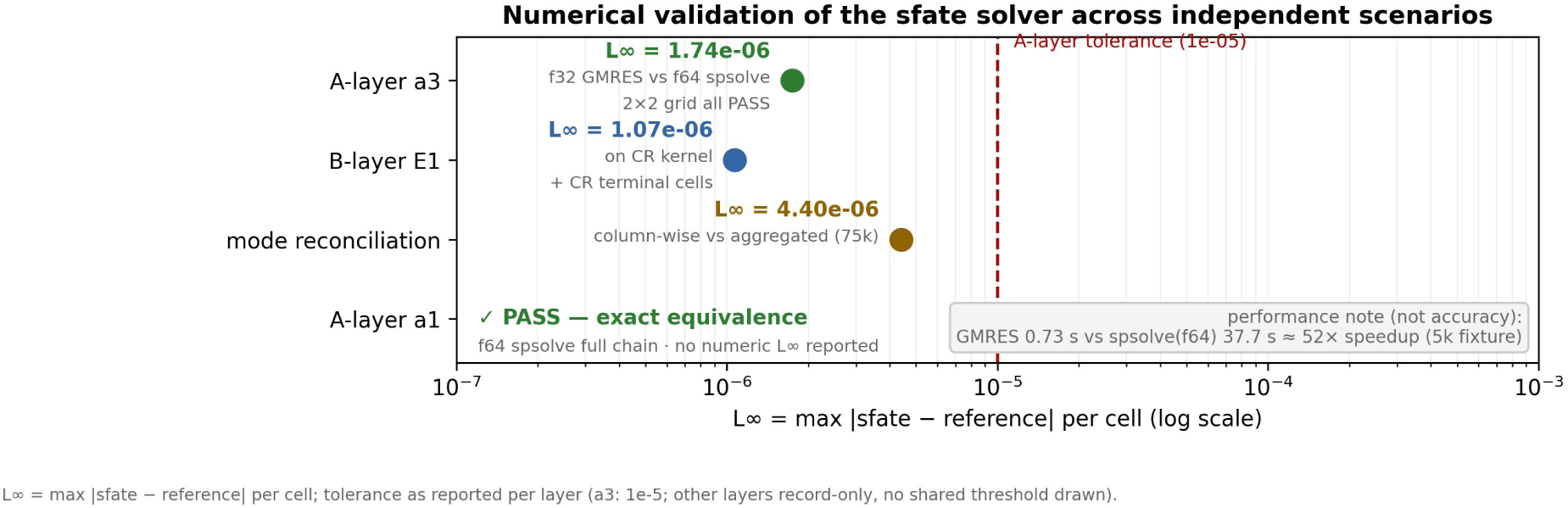
Numerical validation of the sfate solver across independent scenarios (n = 5,000 real-data fixture unless noted). (a) Layer A: production-path agreement with the float64 direct golden reference (all pass L∞ ≤ 1e-5). (b) Layer B: sfate vs CellRank fate probabilities on CellRank’s own kernel and terminal cells (L∞ = 1.07e-06). (c) Reconciliation of aggregated-RHS and column-wise modes (L∞ = 4.4e-06). (d) Timing: float64 direct reference 37.7 s vs production GMRES 0.73 s. Layer C real-data comparisons are record-style and not shown as equivalence tests.

## Results 4 — Spectral terminal-state curation and annotation-defined targets yield different fate landscapes

A first warning appears already at 5k, and we separate the automatic behavior from the comparison setup explicitly. (i) Under fully automatic settings—the eigengap heuristic for the number of macrostates and default terminal-state identification (stability threshold 0.96)—CellRank’s GPCCA pipeline [reuter2018gpcca, reuter2019gpcca] returned a single macrostate and exactly one terminal state (IFN_responsive_1). (ii) Because a six-state comparison was required, ‘n_states = 6’ was then explicitly imposed (a documented override, not an automatic output). (iii) Under that imposed six-state coarse-graining, terminal-state identification produced the fate columns analyzed below. This degeneracy is not a solver artifact: Results 3 established that, on the same kernel and the same terminal cells, sfate’s solver and CellRank’s solver agree to L∞ = 1.07e-06. Whatever drives the discrepancy therefore enters upstream of the linear solve, in the curation step that decides which states count as terminal.

The same behavior recurs at 75k on real data—the two scales are **structurally similar**. In the existing step12 analysis, GPCCA produced five terminal macrostates but split IFN_responsive into three substates (IFN_responsive_1/2/3), declared DAM_like and PU1low_lymphoid **unsupported**, and emitted no fate column for Proliferating despite its macrostate existing. The contingency structure is revealing (Figure 3a): of the 30 DAM_like representative cells, none landed in a macrostate of their own annotation—they scattered across **five** of the six macrostates (8/8/2/4/6/0), and PU1low_lymphoid representatives were nearly absent altogether (5 of 180 total). Representative selection is therefore not incidental: random 30-cell selections (10 seeds) span a median per-state Spearman of 0.450 (5–95% 0.147–0.615) against the centroid reference, indicating that the centroid-nearest construction contributes materially to the boundary condition; sensitivity to alternative selection schemes beyond random sampling was not evaluated.

**Figure 3.**
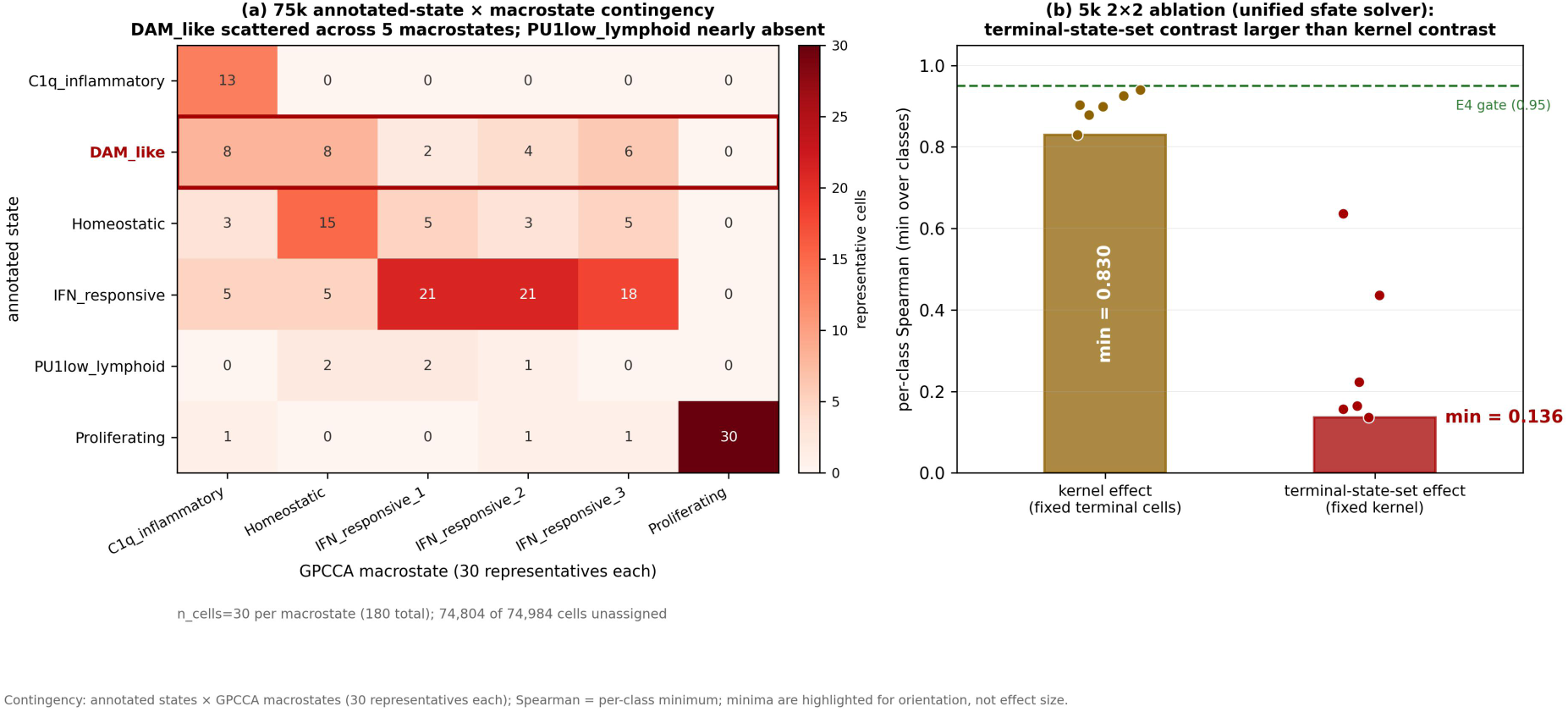
Sensitivity to automatic terminal-state curation. (a) 75k annotated-state × macrostate contingency; numbers are counts of representative cells. (b) 5k 2×2 ablation: per-class Spearman correlations for the kernel contrast (terminal cells fixed; range 0.830–0.940) versus the terminal-set contrast (kernel fixed; range 0.136–0.640). The 0.95 dashed line is a reporting aid, not an acceptance gate. Terminal-set substitution produces the larger observed change, indicating that automatic terminal-state curation is the dominant source of cross-tool divergence in this controlled comparison.

In the 5k 2×2 comparison (kernel × terminal-state set, with a single solver held constant throughout; Figure 3b), terminal-set substitution produced a larger worst-case change than kernel substitution under the settings tested here: the minimum per-class Spearman correlation was **0.136** (per-pair range 0.136– 0.640) for the terminal-set contrast and **0.830** (per-class range 0.830–0.940) for the kernel contrast. Because minima are nonlinear summaries and the six fate columns are not independent, we interpret this as evidence that terminal-state selection is an **important contributor** to the observed cross-tool discrepancy—not as a complete variance decomposition, and not as a causal percentage. The end-to-end disagreement one would naively record between the two pipelines (supplementary-arm minimum Spearman = −0.392) shows that terminal-set substitution produced the larger observed change under the tested controlled comparison, relative to kernel substitution or solver choice; the solver contributed only numerical-tolerance-level differences (L∞ = 1.07e-06). The full six-class paired correlations for both contrasts, with the class-matching rules, are reported in output/b_layer_report.md.

We are careful to delimit what this does and does not mean. GPCCA identifies macrostates from a dominant invariant subspace and is designed to resolve metastable structure [deuflhard2005pcca, roblitz2013pcca, reuter2018gpcca]; PCCA+-type coarse graining finds nearly invariant aggregates, and the annotated states not selected as terminal by the automatic workflow may be less metastable under the chosen kernel—but this interpretation was not tested by an independent metastability analysis in this study, and the literature does not establish that annotation-defined states along transition manifolds are in general excluded by GPCCA-type selection [frank2024spectral]. Consistent with a criterion-driven reading, the automatic terminal-state set is itself threshold-dependent: sweeping the stability threshold over 0.90 / 0.93 / 0.96 / 0.99 on the 5k fixture yields 3 / 2 / 1 / 0 terminal states (Methods; output/threshold_sweep.md)—the curation output is a function of a tunable criterion rather than a stable property of the data. Two further observations constrain the interpretation: the behavior recurs across two scales, and the archived step12 production run is consistent with the krylov-Schur path (a provenance-based inference; Results 1), so the pattern is not unique to our measured dense fallback case. It does not, however, establish that the same pattern must occur under every Schur implementation or parameterization. Our claim is therefore scoped to the curation step, not to the correctness of GPCCA as a method: where annotations exist, an automatic terminal-state decision is an *optional* layer, and its conservative behavior should be reported, not silently inherited. Figure 3 summarizes both lines of evidence; Results 5 shows how annotation-driven target semantics bypasses this curation layer entirely in a real 75k analysis.

## Results 5 — Real-data case: a 75k Alzheimer’s disease mouse microglia fate landscape on a workstation

We applied sfate end-to-end to 74,984 forebrain microglia from a mouse 5xFAD/Cd28-cKO study comprising 12 samples across three genotypes (control, 5xFAD, 5xFAD;Cd28-cKO) at 3, 6, and 8 months. The processed atlas derives from the study of Ayata et al. [ayata2025]; the public source data are available under GEO accession GSE296768. The full pipeline (h5py latent read → approximate kNN graph → six-state absorption solve) ran with a whole-chain peak RSS of **≈1.76 GiB** on the same ∼30 GB workstation on which the default-installation CellRank route back-calculates to ∼405 GB (Results 1). Graph construction took 65.0 s in this first end-to-end run and 18.4 s in the M1 benchmark (≥3 repetitions); the runs differed in execution context, and because cache state and I/O contributions were not isolated experimentally we report both values without assigning the difference to a single cause. The six-state solve itself took **29.0 s** in column-wise mode (∼59 GMRES iterations per column, peak 120; maximum true residual 1.9e-7; zero float64 fallbacks).

Because all six annotated states were supplied as absorbing classes, the output is a complete fate panorama with **no curation step**: every cell carries a probability for every annotated state (Figure 5). We emphasize the semantics of this design choice: designating an annotated class as absorbing defines an **annotation-defined target state**—a user-controlled, auditable modeling decision—not a claim that the class is a biological terminal state (Proliferating, for instance, is included as a target without asserting it is a terminal fate). The number of representatives per class, r, is likewise an explicit parameter, and we measured its impact directly on the 5k fixture under the default column-wise configuration for r = 10/20/30/50/100/200 (reference = r = 200). Rank correlations against the r = 200 reference improve monotonically with r (median Spearman 0.42 → 0.55 → 0.60 → 0.69 → 0.81 → 1.00); per-state mean probability differences fall below 0.034 at r = 30 and below 0.014 at r = 100; and the pointwise L∞ difference remains structurally large at smaller r (≈0.93–0.96) because changing the absorbing set reassigns probability mass at representative cells. A linear mixed model of agreement on log2(r) with state-level random effects gives β = 0.115 per doubling (95% CI 0.098–0.132). r = 30 is therefore reported as an operating point rather than an optimized value.

**Figure 5.**
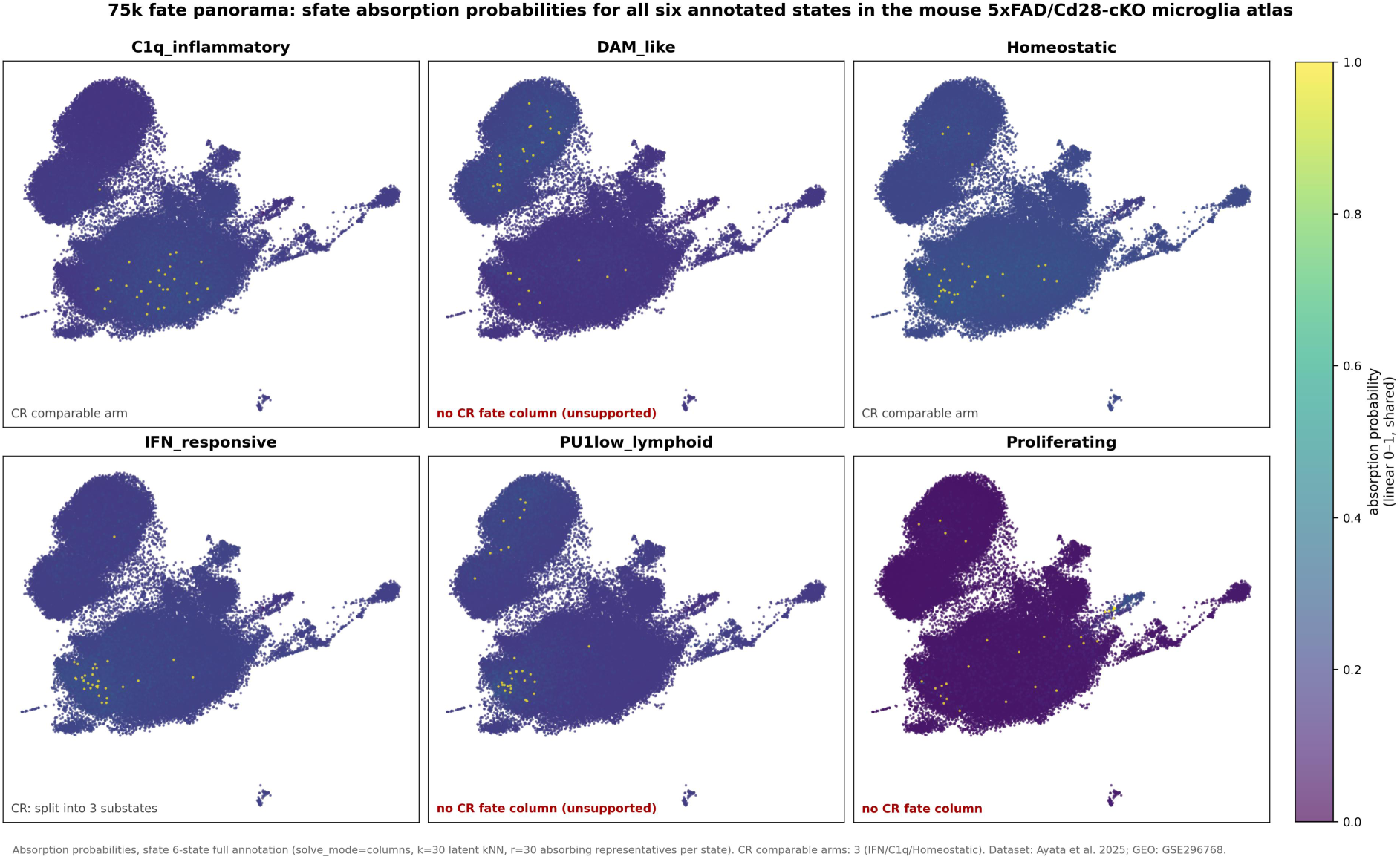
75k fate panorama: sfate absorption probabilities for all six annotated states in the mouse 5xFAD/Cd28-cKO microglia atlas. Each panel shows one target state’s per-cell absorption probability over the UMAP embedding (approximate kNN graph, k = 30; r = 30 absorbing representatives per state, taken as the latent-nearest cells to each class centroid; probabilities normalized across the six supplied target states). Yellow markers denote the absorbing representatives of the corresponding state. Dataset: 74,984 forebrain microglia, 12 samples, three genotypes × three ages [ayata2025; GEO: GSE296768].

The coverage difference against the existing step12 CellRank analysis is factual and structural: the CellRank side offers only three comparable fate arms (IFN_responsive, C1q_inflammatory, Homeostatic), while DAM_like, PU1low_lymphoid, and Proliferating have no fate columns at all—DAM_like and PU1low_lymphoid were returned as "unsupported" by the curation step, and Proliferating produced no output column. The missing arms are not peripheral states: disease-associated microglia (DAM) are a disease-restricting microglial type first defined in Alzheimer’s models [kerenshaul2017dam] and a proposed universal immune sensor of neurodegeneration [deczkowska2018dam], positioned within the homeostatic-to-neurodegenerative microglial program [krasemann2017], and the IFN-responsive/C1q-inflammatory arms intersect with the type-I-interferon–driven neuroinflammatory axis described in Alzheimer’s disease [roy2020ifn]. We emphasize that this is a statement about output coverage, not about which tool is "right" about disease mechanism: the two pipelines answer differently-scoped questions, and the divergence is attributable to the curation layer analyzed in Results 4, not to stochastic error.

Where the two sides can be compared, the disagreement is structured rather than diffuse: it is concentrated in specific fate arms instead of being spread uniformly. Structured here refers to that fate-specific concentration; overall direction concordance itself did not exceed sign randomization (p_all = 0.339). At sample level (12-sample aggregation; genotype-delta sign over a 3 fates × 2 contrasts × 3 ages grid = 18 comparison cells, of which 3 are not comparable because no cKO sample exists at 8 months, leaving 15 scored comparison cells), the two pipelines agree on the genotype-delta direction in **9 of 15 comparison cells** (Figure 6a)—we stress that these are 15 contrast × fate × age cells sharing samples, not 15 independent observations, so no binomial-style significance claim attaches to the proportion. The breakdown is fate-specific rather than globally concordant: on the 5xFAD−control contrast the agreement is mixed (5/9); on the cKO−5xFAD contrast it is **4/6** in total, with complete directional agreement for IFN_responsive and C1q_inflammatory (4/4) and complete reversal for Homeostatic (0/2). This Homeostatic-specific reversal resonates with two independent observations: at the single-cell level, Homeostatic is precisely the arm where the two sides anti-correlate (per-cell Spearman: IFN_responsive 0.320, C1q_inflammatory 0.328, **Homeostatic −0.568**, versus a CellRank commitment score of 0.916 on its own three arms), and in the Results 4 ablation the weakest terminal-state mapping pairs involve the Homeostatic macrostates (Homeostatic_1→IFN_responsive 0.136; Homeostatic_2→Homeostatic 0.223). These concordant summaries identify where the workflows differ, but they do not determine which fate estimate is biologically more accurate; independent lineage information or perturbational validation would be required for that conclusion. A sign-permutation test over the 15 scored comparison cells yields p_all = 0.339 for the overall 9/15 agreement; per-fate agreement is 4/5 for both IFN_responsive and C1q_inflammatory (p = 0.188 each), and p = 0.969 against the complete-reversal alternative for Homeostatic (the observed opposition of mean directions does not exceed the sign-randomization expectation conditional on the marginal directions). These null distributions do not support a significant shared directional signal, so we retain the descriptive interpretation above. The pattern is consistent with the terminal-state mapping differences identified in the controlled ablation; it does not by itself isolate GPCCA’s mapping as the sole cause, given the remaining differences in kernels, representative cells, and normalization between the two pipelines.

**Figure 6.**
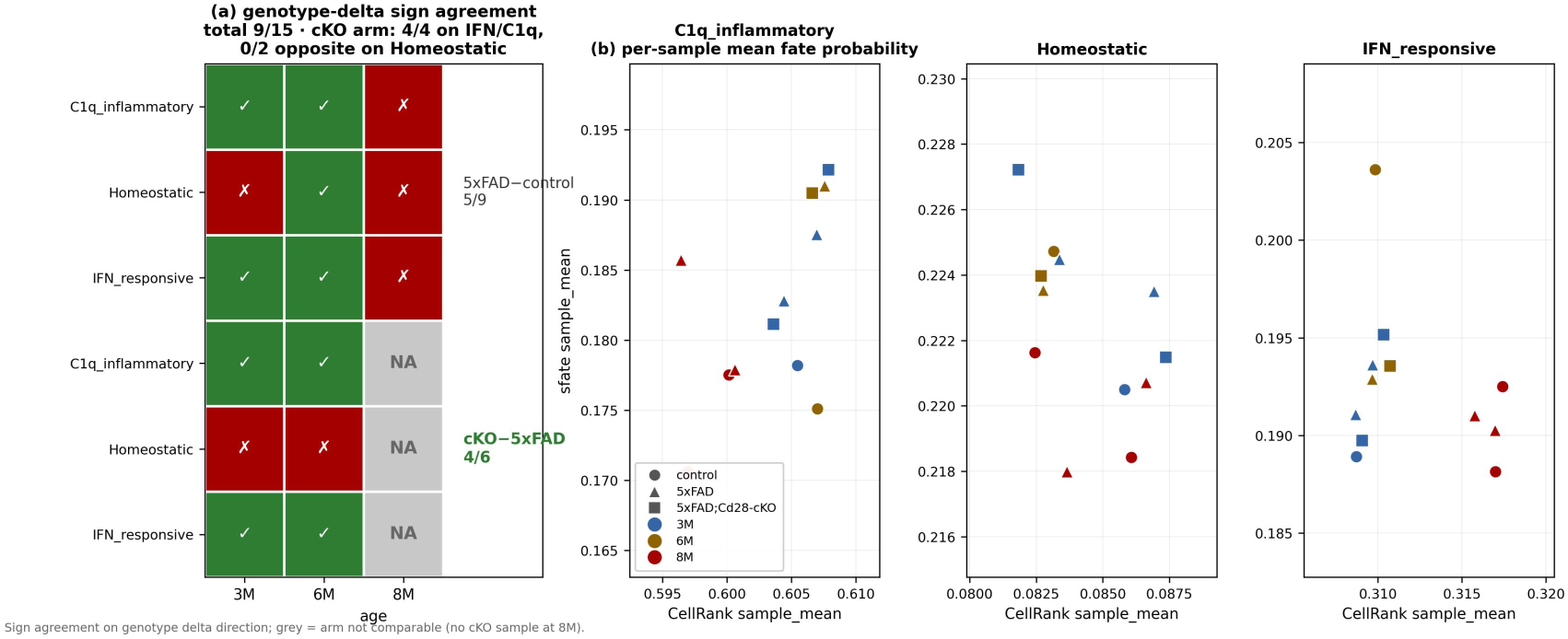
Sample-level direction consistency between sfate and the archived CellRank analysis. (a) Genotype-delta sign-agreement grid over 3 fates × 2 contrasts × 3 ages = 18 cells, of which 3 are NA (no cKO sample at 8 months), leaving 15 scored cells; overall agreement 9/15 (5xFAD−control 5/9; cKO−5xFAD 4/6, with IFN_responsive and C1q_inflammatory 4/4 and Homeostatic 0/2, opposite direction). Delta direction is defined as (mutant − reference) mean fate probability per sample. (b) Per-sample mean fate probabilities by genotype and age; points are samples, not independent observations. Direction agreement is assessed by the sign-permutation test in the text (Results 5); this panel itself is descriptive; the 15 scored cells share samples and are not independent.

The 75k run also produced a methodological finding about the solve itself (Figure 4). In the initial aggregated-RHS configuration (class-summed right-hand sides, ILU(1) preconditioning), 5 of 6 classes failed to reach rtol = 1e-6 within 50,000 GMRES iterations and the solve cost 3,915 s including float64 fallbacks. The final configuration—column-wise solves (one per absorbing cell, class aggregation after the solve; mathematically equivalent by linearity, as certified by the a1 lumping check in Results 3) without a preconditioner—converged rapidly: ∼59 iterations per column and **29.0 s** for all six classes, a 135× difference from the initial configuration (3,915 / 29.0 ≈ 135.0). Because both the right-hand-side organization and the preconditioner changed between these two configurations, this comparison does not identify either factor as the sole cause; the observed convergence difference is consistent with stronger projection of aggregated right-hand-side vectors onto slowly decaying modes of Q, although GMRES convergence for this generally non-normal system cannot be predicted from eigenvalues alone [titleypeloquin2014, sharpewales2021]. The separate ILU build measurements—spilu construction 132 s at fill = 1 and 380 s at fill = 10, versus 0.1 s for the reported unpreconditioned single-column solve— show that these ILU settings were not cost-effective for this matrix; the comparison is between ILU factorization build time and a single unpreconditioned column solve, not between end-to-end configuration totals [saad1994ilut, benzi2002survey].

**Figure 4.**
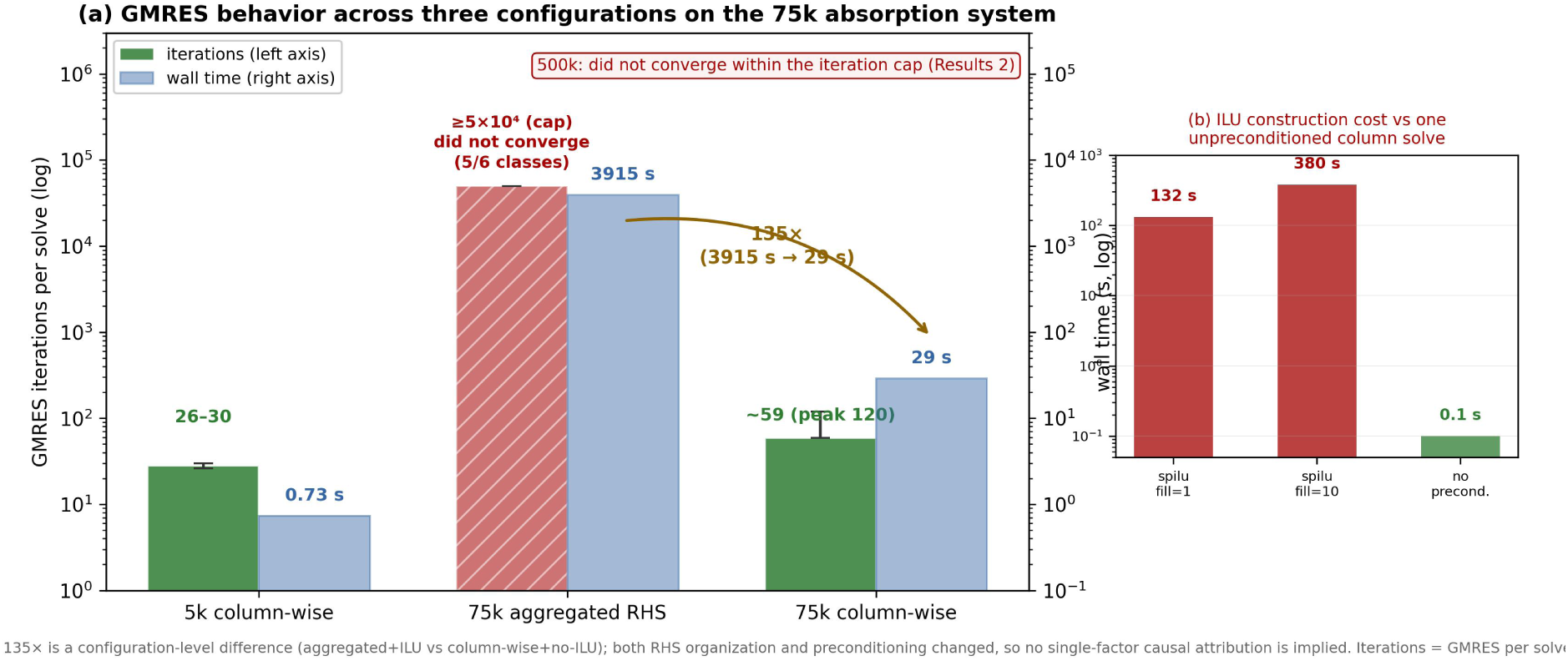
GMRES behavior across three configurations on the 75k absorption system (n = 74,984; six annotated absorbing classes; rtol = 1e-6; restart = 50; at most 1,000 restarts). Curves show true-residual history per configuration: aggregated-RHS + ILU(1) (5 of 6 classes failed to reach rtol = 1e-6 within 50,000 iterations; 3,915 s including float64 fallbacks) and column-wise without preconditioner (∼59 iterations per column; 29.0 s total). The 135× figure is a configuration-level difference between aggregated+ILU and column-wise+no-ILU: both right-hand-side organization and preconditioning changed, so no single-factor causal attribution is implied. Inset: ILU factorization build cost (132 s at fill = 1; 380 s at fill = 10) versus 0.1 s for one unpreconditioned column solve; these are different operations and the inset is not an end-to-end comparison.

In summary, on real 75k data sfate delivers six annotation-defined fate columns, whereas the archived automatic GPCCA workflow emitted three comparable fate columns under the settings used here (Table 2); GPCCA-based pipelines can also accept prior knowledge or user-specified state choices, so the comparison concerns the observed automatic-curation output, not a structural impossibility of the estimator. The complete panorama was obtained at ≈1.76 GiB peak memory on consumer hardware. The two workflows show structured differences in the fate arms returned under the tested configurations: the disagreement is concentrated in specific arms, sign-structured across scales, and consistent with the behavior of the automatic curation layer that sfate replaces with explicit, annotation-controlled target semantics.

**Table 2.** Cross-tool consistency evidence (Layers B and C)

| Layer/comparison | Metric | Value | Scope and interpretation |
| --- | --- | --- | --- |
| B-layer · E1 cross-validation | $L_\infty$ | 1.07e-06 | [MEASURED] CR kernel + CR terminal cells; sfate vs CellRank fate; solver agreement at tested tolerance |
| B-layer · kernel contrast (terminal cells fixed) | min Spearman | 0.830 | [MEASURED] per-class 0.830–0.940; kernel substitution is smaller contrast |
| B-layer · terminal-set contrast (kernel fixed) | min Spearman | 0.136 | [MEASURED] per-pair 0.136–0.640; terminal-set substitution is larger contrast |
| B-layer · end-to-end supplementary arm | min Spearman | -0.3918 | [MEASURED] red-flag disagreement; not a decision criterion (main arm degenerated to one terminal state) |
| C-layer · per-cell consistency (3 comparable arms, 75k) | Spearman | IFN 0.320 / C1q 0.328 / Homeostatic -0.568 | [MEASURED] record-style comparison, no gate; low correlation expected because terminal-state sets differ by construction; CR commitment_score mean 0.916 |
| C-layer · sample-level direction consistency (12-sample aggregation) | sign agreement | 9/15 | [MEASURED] 5xFAD–control 5/9; cKO–5xFAD 4/6 (IFN 2/2 + C1q 2/2 + Homeostatic 0/2); 15 = 3 arms × 2 contrasts × 3 ages – 3 NA (cKO 8M no sample) |

## Discussion

Our results position sfate within an engineering tradeoff rather than a performance race. On one pole stands the **heavy-dependency** route: CellRank with a correctly provisioned PETSc/SLEPc/MPI stack [balay2026petsc, hernandez2005slepc], which—as our own measurements in Results 1 confirm— eliminates the quadratic memory wall at least to the half-million-cell scale. On the other pole stands the **light-dependency** route sfate takes: pure scipy, installable by ‘pip’ alone. sfate removes the specific Schur fallback examined here; this design does not establish the absence of all scale-dependent failure modes elsewhere in the pipeline. The wall we documented is therefore scoped precisely: it is the default installation’s Brandts fallback, not a ceiling on what GPCCA can compute. We do not read this as a criticism of SLEPc—an effective, mature eigensolver—but as a statement about what a user gets when the optional stack is absent, mis-provisioned, or simply unavailable in a restricted environment. PETSc/SLEPc are optional in the examined CellRank installation path and may require source compilation under pip, whereas conda-forge provides pre-built packages on supported platforms [cellrankinstallation]. The subject of this tradeoff is the dependency stack’s engineering properties, not any community’s choices.

Crucially, sfate’s value proposition does not collapse in the krylov-available regime; three of its pillars are orthogonal to the memory wall entirely. First, the **explicit validation protocol** (Results 3): numerical agreement with a float64 direct reference and with CellRank’s own solver output on identical inputs, within a prespecified tolerance [higham2002accuracy]. Second, **annotation-controlled target semantics** (Results 4): where cell-type annotations exist, absorption probabilities can be computed for every annotation-defined target state without an automatic curation step. (CellRank does expose manual terminal-state override, but the workflow remains GPCCA-centered: the Schur step stays in the estimator path, the PETSc/SLEPc dependency remains for the sparse route, and no numerical certificate is produced; sfate instead exposes the absorption problem as the primary computational object.) Third, the kernel/terminal-set/solver decomposition itself (Results 4): the conservative behavior of GPCCA’s terminal-state curation—observed in the archived krylov-Schur output—is a methodological observation about metastability-based coarse-graining that stands regardless of which Schur route is used. These are **properties of the present implementation and evaluation design**; we do not claim that other software stacks cannot provide equivalent validation or user-specified terminal states.

The aggregate-after-solve observation generalizes cautiously beyond sfate. Linearity guarantees that per-representative solutions may be summed after solving, but it does not guarantee that this organization will converge faster for every matrix or right-hand side: right-hand-side-dependent GMRES convergence is well documented [titleypeloquin2014], and multi-right-hand-side Krylov methods are an active topic [simoncini1996block, sukmanyuk2024]. In our 75k run, the column-wise, unpreconditioned configuration was markedly faster than the aggregated, ILU-preconditioned configuration. The result motivates testing aggregate-after-solve in other absorption-probability implementations, while the causal roles of right-hand-side organization, restarting, warm starts, and ILU require a factorial comparison, which we leave as stated future work. Likewise, the observed ILU overhead is restricted to the matrix and ‘spilù settings reported here (geometric kNN transition graphs with hub structure); a hub-mediated failure mechanism was not independently established, and we do not claim the observation generalizes to other preconditioning contexts [benzi2002survey, chowsaad1997].

Six limitations deserve explicit statement. First, the million-cell regime is supported only by extrapolation of the measured four-tier linear fit [EXTRAPOLATED]; the fit’s slope is dominated by kNN-index scratch allocations, which are optimizable but not yet optimized. Second, the 500k tier now covers graph construction only as a successful measurement; a full-pipeline 500k attempt under the default column-wise GMRES configuration did not converge for the first absorbing cell within 49,904 iterations (true residual 1.09e-5 at rtol = 1e-6). This is a scale-dependent solver observation, not a ceiling on all possible configurations: restarting, preconditioning, right-hand-side organization, or warm starts may recover convergence, and a factorial comparison is required before any single factor is identified as the cause. Until then, the 1M memory extrapolation applies only to graph construction and I/O, and the Figure 1 comparison remains on a graph-construction basis at 500k. Third, our measurements span two CellRank versions: **cellrank 2.1.0** (local environment: B-layer ablation and the krylov-Schur benchmarks) versus **cellrank 2.3.2** (the survey baseline in which the empirical n² model was calibrated); the Brandts fallback measurements were reproduced across both versions to within sampling precision (Methods), and GPCCA defaults and fate-normalization behavior follow 2.1.0 where the two differ, with version-specific differences enumerated in Supplementary Table S1. Fourth, neighbor recall and seed sensitivity were evaluated only on the tractable tiers (5k real: 0.9803 ± 0.0001 across three seeds; 10k synthetic: 0.9945); the accuracy of the approximate graph at 500k, where exact search is intractable, remains unevaluated. Fifth, the 135× configuration comparison confounds right-hand-side organization with preconditioning; a factorial RHS × ILU comparison is required before either factor can be credited individually. Sixth, the absorption landscape is conditional on the supplied embedding: replacing the scVI latent with a PCA-30D embedding of the same cells materially changed the landscape (median per-state Spearman 0.165, range −0.37 to 0.64), and sfate does not infer a representation-independent fate landscape.

Future work proceeds in three directions. First, the factorial comparison isolating right-hand-side organization, preconditioning, restarting, and warm starts on geometric kNN absorption systems, together with spectral diagnostics (right-hand-side projections onto slow modes; residual histories) to test the slow-mode hypothesis. Production hardening will add a directed-reachability (ρ(Q)<1) pre-check so that the stated M-matrix condition becomes a guaranteed rather than a posteriori-verified precondition. Second, for datasets exceeding the linear regime or lacking per-cell resolution, a waypoint-based alternative— conceptually related to Palantir [setty2019palantir]—could reduce the state space before absorption-probability estimation, followed by an explicitly validated mapping back to cells; this would be a **distinct approximation strategy** rather than a degraded form of the present method. Third, accuracy validation of the approximate graph at 500k, where exact search is intractable, and the downstream gene-trend machinery (per-gene × lineage models), which neither CellRank 2’s sparsity improvements nor sfate currently addresses, is the next memory amplifier worth eliminating.

We close with the motivation that framed this study. Single-cell analysis increasingly reaches scales at which a back-calculated 405 GB requirement is inaccessible on a typical workstation. Restricted compute environments may also arise from institutional governance policies, although the mouse dataset analyzed here does not itself test that use case. sfate addresses the local-compute constraint by maintaining linear measured memory over the tested graph-construction tiers and by returning all six requested annotation-defined fate columns at 75k, with numerical agreement assessed under the stated validation protocol.

## Methods

### Data

The study dataset is a processed 74,984-cell atlas of forebrain microglia from the mouse 5xFAD/Cd28-cKO study of Ayata et al. [ayata2025] (12 samples; three genotypes—control, 5xFAD, 5xFAD;Cd28-cKO; three age groups—3, 6, 8 months). Raw and source-level data are available through GEO accession GSE296768. The present analysis used the processed derivative ‘adata_annotated.h5ad’ (74,984 × 55,401) generated by the originating pipeline and stored read-only on institutional infrastructure. Per-cell inputs to sfate are the 30-dimensional scVI latent representation (‘obsm[’X_scVI’]’, (74,984 × 30) float32) and the six-class ‘microglia_statè annotation (Homeostatic 27,953; IFN_responsive 20,303; C1q_inflammatory 13,575; DAM_like 6,796; PU1low_lymphoid 5,560; Proliferating 797). Latent and metadata were read directly with h5py in read-only mode; anndata 0.12.16 backed mode was avoided after measuring that it eagerly materializes all layers (+6.9 GB on opening our annotated h5ad for this file, version, and access path). UMAP coordinates (‘obsm[’X_umap’]’, (74,984 × 2)) were used only for visualization. The reference CellRank products are the existing step12 outputs, used as a **historical reference analysis** (an existing CellRank baseline), not a same-conditions benchmark: the two sides differ in environment, gene representation, and kernel construction by design, and all cross-tool comparisons are interpreted accordingly (Layers B/C). The deposited code and data manifest will map this derivative to the public accession and document each preprocessing stage. All algorithmic benchmarks (Layers A/B and the scaling suite) are fully reproducible from the synthetic fixtures without access to the mouse atlas.

### The sfate pipeline

sfate computes fate probabilities in five parameterized stages. (1) **Latent input**: a low-dimensional representation (here 30-d scVI [lopez2018scvi]; the interface requires (n, d) with d ≤ 256). (2) **Approximate kNN graph**: NN-Descent (pynndescent 0.6.0 [dong2011nndescent], ‘n_neighbors=30’, ‘metric=’euclidean’’, ‘random_state=20260909’; ‘n_trees’, ‘n_iters’, diversification/pruning, and thread count use pynndescent defaults and are reported in Supplementary Table S2). The sklearn exact tree backend is retained as a cross-check branch. (3) **Transition matrix**: kNN edges weighted by a Gaussian kernel, max-symmetrized, row-normalized to a single CSR float32 row-stochastic matrix (nnz ≈ 1.6– 1.7·n·k). The Gaussian bandwidth for row i is the distance to its k-th nearest neighbor, d_k(i); the weight of edge (i, j) is exp(−(d_ij / d_k(i))²), with duplicate/tie handling and row-normalization in float64 followed by a cast to float32. No second copy, no float64 upcast of the full matrix, and no logits are used. The resulting operator is a **diffusion-type row-stochastic matrix on latent-space proximity and contains no velocity-direction term**; the quantities computed from it are absorption probabilities toward user-defined target states on this graph, not velocity-derived lineage fate probabilities. (4) **Absorbing states**: each annotated class contributes its r cells nearest to the class centroid in latent space as absorbing representatives (we used r = 30 as the default representative-set size to match the scale of the historical CellRank analysis, and treated r as an explicit sensitivity parameter; Results 5). The centroid and nearest-cell distances use Euclidean metric in latent space; ties are broken by NumPy’s stable argsort. If a class has fewer than r cells, all cells are used; overlapping representatives across classes are allowed and resolved by the absorbing-state construction. (5) **Absorption solve**: the linear system (I−Q)X = R is solved column-wise with unpreconditioned GMRES, where Q is the transient×transient block and R the transient×absorbing block of the row-stochastic transition matrix after absorbing states are set to unit rows. A directed-reachability/ρ(Q)<1 check is **not implemented** in the current code; the solver guards only against undirected connected components that contain no absorbing cell (‘scipy.sparse.csgraph.connected_components’ with ‘directed=Falsè). All reported validation systems satisfied nonsingular M-matrix assumptions empirically, but this is stated as a validated condition rather than a guaranteed precondition for arbitrary input. Production acceptance requires a recomputed unpreconditioned true residual below the solver tolerance; probabilistic invariants (row-stochasticity, one-hot absorbing rows, nonnegativity of F) hold by construction and are exercised in the test suite.

### Solver modes and defaults

Three modes are implemented and were used for distinct purposes in this paper. The **default** is ‘solve_mode="columns"’ (per-absorbing-cell solves with post-hoc class aggregation, exploiting linearity; no preconditioner). ‘solve_mode="aggregated"’ (class-summed right-hand sides, ILU optional) is retained for reconciliation against the default (the two modes agree to L∞ = 4.4e-06; Results 3). SciPy 1.15.3 ‘gmres’ is called with ‘rtol=1e-6’, ‘atol=0.0’, ‘restart=50’, ‘maxiter=1000’, and ‘callback_type=’pr_norm’’; convergence is accepted only after recomputing the unpreconditioned true residual in the working dtype (float32 for production, float64 for the strict validation path). A float64 direct solve (scipy ‘spsolvè) serves exclusively as the small-scale golden standard for validation, never as a production path (fill-in explosion at 5k already costs 37.7 s per class). If a column fails the production criterion, the fallback recomputes the system in float64 (A, RHS, and solution all float64) and reuses the float32 ILU factors as a mixed-precision preconditioner if ILU was requested. ‘block_size=256’ denotes the number of RHS columns processed together in the outer loop over targets; it bounds peak memory by controlling the size of the dense solution buffer and does not change the mathematical system, as verified by the reported block-size sweep.

### Validation protocol

Acceptance uses a three-layer protocol with explicit references. **Layer A**: on identical transition matrices and absorbing sets, the production path (float32 GMRES) is compared against the float64 ‘spsolvè golden reference. Acceptance combines two criteria with distinct roles: (i) a **strict solver-level criterion**—the computed iterate must have a normalized backward error ≤ 1e-12, indicating that the computed iterate is consistent with the exact solution of a nearby linear system within the stated backward-error bound (applied in the a2 unit test, n = 1,600 orphan-free synthetic chain, both solve modes, GMRES tol = 1e-12); and (ii) a **lenient application-level tolerance**—observed L∞ ≤ 1e-5 against the float64 reference (applied to the production path, tol = 1e-6, 5k fixture, a3). Because no directed-reachability check is implemented in the production path (see The sfate pipeline), reachability and the nonsingular M-matrix property were verified a posteriori on each validation fixture by exact computation of A⁻¹·1 in float64; for arbitrary input these remain stated conditions rather than guaranteed preconditions. Under these conditions, A⁻¹ ≥ 0, so ‖A⁻¹‖∞ = max(A⁻¹·1) and κ∞(A) = ‖A‖∞·max(A⁻¹·1). The reported normwise backward error is η = ‖Ax−b‖∞ / (‖A‖∞‖x‖∞ + ‖b‖∞), computed in float64 for the strict tier and float32 for the production tier; the forward discrepancy is absolute L∞ = max|x − x_ref|. The strict AND-criterion (backward error ≤ 1e-12 and L∞ ≤ 10·κ∞·backward_max) applies to the float64 validation tier (a2). The float32 production path (a3) cannot attain 1e-12 backward error and is gated by the application-level tolerance L∞ ≤ 1e-5 alone. The two tiers are not interchangeable thresholds but different statements: the strict tier certifies the solver, the lenient tier certifies fitness for biological interpretation. The ∼400× norm-estimator understatement refers to a comparison between ‘scipy.sparse.linalg.onenormest’ and the exact κ∞ computed from the M-matrix identity on the 5k fixture; the exact identity is therefore used in place of the estimator.

The reconciled four-value certificate (regenerated 2026-09-10 by ‘scripts/cert_recompute_5k.py’, full log in ‘output/cert_recompute_5k.md’):

| Tier | backward error | $\kappa_\infty(I-Q)$ | $10 \cdot \kappa_\infty \cdot \text{backward}$ | observed $L_\infty$ | gate | PASS |
| --- | --- | --- | --- | --- | --- | --- |
| Strict a2 (aggregated, f64 tol=$1e-12$) | 1.35e-13 | 5.01e+02 | 6.78e-10 | 8.78e-13 | $1e-12$ | ✓ |
| Strict a2 (columns, f64 tol=$1e-12$) | 2.08e-13 | 5.01e+02 | 1.04e-09 | 9.28e-13 | $1e-12$ | ✓ |
| Production a3 (columns+none, current default) | 3.27e-07 (normalized $\eta$) | 3.21e+01 | 1.05e-04 | 1.74e-06 ($5.7\times$ vs $1e-5$) | $1e-5$ | ✓ |

**Layer B**: the sfate solver is run on CellRank’s own kernel and terminal cells and reconciled against CellRank fate probabilities (cross-validation, E1). **Layer C**: full-data comparison against existing step12 products, record-only with no acceptance gate (terminal-state sets differ by construction).

### Benchmarking protocol

All scaling numbers were measured single-process with BLAS threads = 1 and fixed seed 20260909, at ≥3 repetitions per tier (maximum reported). Benchmarks were run on a consumer workstation (Mechrevo JIAOLONG 16; AMD Ryzen 9 8945HX, 16 cores / 32 threads, boost up to 5.46 GHz), with 32 GB of installed RAM (1× 32 GB DDR5-5600 SODIMM, Crucial CT32G56C46S5.C8BA; 30 GiB OS-visible, the remainder hardware-reserved), on Ubuntu 24.04.4 LTS (kernel 7.0.0-31-generic), with the working directory on an internal YMTC PC41Q 1 TB NVMe SSD (ext4). The system had 46 GB of configured swap (15.6 GB swapfile + 30.5 GB partition); /usr/bin/time -v reported Swaps: 0 in every reported run. We set and report OMP_NUM_THREADS = 1 and OPENBLAS_NUM_THREADS = 1; the benchmark environment (Python 3.12, conda env rfd3) links scipy-openblas 0.3.31.dev via SciPy 1.15.3 / numpy 2.4.2 (MKL not present). CPU frequency scaling remained enabled, as is typical for consumer workstations. For each tier, Supplementary Table S3 gives the exact number of repetitions, warm-up policy, cache state, process affinity, and whether the reported maximum combines separate runs. RSS was sampled every 250 ms and cross-checked against ‘/usr/bin/time -v’; the sampler’s own memory and child-process handling are documented. The synthetic generator, all parameters, and output checksums are provided in the repository.

The 500k full-pipeline benchmark was attempted under identical single-process settings. Graph construction completed in 66.2 s with a whole-process peak RSS of 4.03 GiB, but the default column-wise GMRES solve did not converge for the first absorbing cell within 49,904 iterations (true residual 1.09e-5 at rtol = 1e-6). The 500k absorption solve is therefore reported as a negative result under the default configuration; the 1M memory extrapolation applies only to graph construction and I/O, not to the absorption stage.

### Sensitivity and robustness analyses

Representative-count sensitivity was assessed on the 5k real fixture for r ∈ {10, 20, 30, 50, 100, 200} against a reference at r = 200, using the default column-wise solver. Agreement was summarized per absorbing state by Spearman correlation of cell-wise fate probabilities and by mean absolute probability difference; a linear mixed model with log2(r) as fixed effect and state as random intercept was fit with ‘statsmodels.mixedlm’ to obtain the per-doubling improvement β and 95% confidence interval. kNN-neighbor sensitivity used the same fixture with k ∈ {15, 30, 50, 100} and r = 30, reporting Spearman correlation and MAE relative to k = 30 and graph-construction time. Latent-representation sensitivity compared the scVI-30D reference to a PCA-30D embedding of the same cells (top 2,000 highly variable genes, sklearn PCA, fixed seed) under the identical k = 30, r = 30 production protocol, reporting per-state Spearman correlation and MAE. Representative-selection sensitivity fixed the kNN graph and compared the production centroid-nearest r = 30 selection to 10 random within-class 30-cell selections (10 seeds) on the 5k fixture, reporting per-state Spearman correlation and MAE against the centroid reference and the random-selection overlap rate.

Sample-level directional agreement between sfate and CellRank was tested by a sign-permutation procedure. Each of 15 comparable comparison cells (3 fates × 2 genotype contrasts × 3 ages, minus 3 cells with no 8-month cKO sample) records the sign of the genotype-delta for sfate and CellRank. The overall null distribution was generated by 10,000 random flips of one pipeline’s sign vector across all 15 cells; the per-fate nulls used exact enumeration over the 5 cells per fate. The Homeostatic reversal was tested against the specific alternative of complete sign reversal (one-tailed exact p-value).

### Environment and reproducibility

Environment fingerprint: Python 3.12.9; cellrank 2.1.0; pygpcca 1.0.4; scanpy 1.11.5; anndata 0.12.16; scipy 1.15.3; numpy 2.4.2; pandas 2.3.3; h5py 3.16.0; pynndescent 0.6.0; petsc4py 3.25.5 and slepc4py 3.25.1 (PETSc lib 3.25.5) for the krylov-Schur control only; glibc 2.39; Linux x86_64; ∼30 GiB physical RAM. **Version deviation**: the survey baseline on which the empirical n² model was calibrated is cellrank 2.3.2 (sandbox), reproduced to within sampling precision on cellrank 2.1.0 (Brandts 2k/5k points, output/brandts_8k.md); all local experiments (Layer B, Layer C, krylov benchmarks) ran on cellrank 2.1.0; GPCCA defaults and fate-normalization behavior follow 2.1.0 where the two differ, with version-specific differences enumerated in Supplementary Table S1. Software citations follow first use: CellRank [lange2022cellrank, weiler2024cellrank2], GPCCA [reuter2018gpcca, reuter2019gpcca], scVI [lopez2018scvi], NN-Descent [dong2011nndescent], GMRES [saad1986gmres], scanpy [wolf2018scanpy], PETSc/SLEPc [balay2026petsc, hernandez2005slepc], brandts Schur [brandts2002schur, anderson1999lapack]. **Code availability**: sfate source, validation suite (32 unit tests, deterministic under fixed seed), benchmark scripts, and figure-generation scripts are archived at https://github.com/Ericleo-zeng/sfate (release tag v1.0.0, commit 5472f62714285ebcfae03b54f4c387de0fdbf69a, MIT license) and archived at Zenodo (DOI: 10.5281/zenodo.22851715). The package includes the 5k validation fixture and the 75k fate matrices. The scVI latent matrix and annotation table accompany the preprint deposition; raw and source-level data are public (GEO: GSE296768).

**Units.** GB denotes 10⁹ bytes; GiB denotes 2³⁰ bytes. Rounded values are used only in the title and narrative shorthand; tables report recorded precision. "75k" is shorthand for 74,984 cells.

## Footnotes

1 CellRank issue #540, "Scaling CellRank to large datasets," maintainer comment: https://github.com/scverse/cellrank/issues/540#issuecomment-804096208

2 CellRank issue #1146, materialization of a moscot transport coupling: https://github.com/scverse/cellrank/issues/1146#issuecomment-1853492566

## Supplementary Tables

**Table S1.** Version differences between CellRank 2.1.0 and 2.3.2

| Component | 2.1.0 (local experiments) | 2.3.2 (survey baseline) | Relevance to this study |
| --- | --- | --- | --- |
| Schur fallback when PETSc/SLEPc absent | ``method='brandts'`` after warning; densifies sparse input | Same warning + ``brandts`` densification (commit 4719939) | Empirical $n^2$ model calibrated in 2.3.2 sandbox and reproduced on 2.1.0 |
| Schur method selection API | ``compute_schur(method='krylov'\|'brandts')`` | Same | — |
| Macrostates: ``cluster_key`` | ``str`` only (obs categorical) | ``str`` \| Mapping supporting ``obs`` or ``obsm`` proportions; added ``weight_key`` | Krylov-Schur controls run under 2.1.0 |
| Terminal-state selection: stability method | No explicit error when no state meets threshold | Raises ``ValueError`` if ``stability_threshold`` too high | Not used; no macrostate computation in sfate; 5k fixture behavior observed under 2.1.0; auto selection still falls back to ``n_states=6`` when eigengap degenerates |
| ``set_terminal_states`` aggregation | ``top_n`` only | Added ``agg`` parameter (``top_n`` / ``union``) | Not used; representative-cell scheme in sfate is explicit |
| Fate-probability normalization | Row-normalized absorption classes; ``check_sum_tol=1e-3`` | Same core computation; plot method gained ``basis`` argument | Layer B/C comparisons use 2.1.0 outputs; normalization consistent across versions |
| Logging | Custom ``cellrank.logging`` | Standard ``logging`` module | No numerical impact |

Notes: Differences are primarily API conveniences and error handling; the Schur/Brandts fallback path and fate-probability core are unchanged between the two versions for the settings used here.

**Table S2.** Pipeline parameter defaults

| Stage | Parameter | Default value | Notes |
| --- | --- | --- | --- |
| Latent input | ``d`` | $\leq 256$ | Interface requires a dense 2-D array; this study uses 30-d scVI |
| kNN graph | backend | ``pynndescent`` | ``n_neighbors=30``, ``metric='euclidean'``, ``random_state=20260909`` |
| kNN graph | ``n_trees`` | ``None`` (pynndescent default) | pynndescent selects automatically |
| kNN graph | ``n_iters`` | ``None`` (pynndescent default) | pynndescent selects automatically |
| kNN graph | ``diversify_prob`` | 1.0 | pynndescent default |
| kNN graph | ``pruning_degree_multiplier`` | 1.5 | pynndescent default |
| kNN graph | ``leaf_size`` | ``None`` | pynndescent default |
| kNN graph | ``n_jobs`` | ``None`` | Runs single-threaded in this study |
| Transition matrix | kernel | Gaussian | $w_{ij} = \exp(-(d_{ij} / d_{k(i)})^2)$, where $d_{k(i)}$ is the distance to the $k$-th nearest neighbor |
| Transition matrix | symmetrize | ``max`` | Max-symmetrized before row normalization |
| Transition matrix | dtype | float32 | Row-normalized in float64, then cast to float32 |
| Absorbing states | ``r`` | 30 | Centroid-nearest cells per annotated class in latent Euclidean space; ties broken by NumPy stable argsort |
| Absorbing states | centroid metric | Euclidean | Computed in the same latent space used for the kNN graph |
| Absorption solve | solver | SciPy 1.15.3 ``gmres`` | ``rtol=1e-6``, ``atol=0.0``, |
|  |  |  | ``restart=50`, `maxiter=1000`,`<br>``callback_type='pr_norm'`` |
| Absorption solve | solve mode | ``columns`` | Per-absorbing-cell solves,<br>class aggregation after solving |
| Absorption solve | preconditioner | ``none`` | ILU optional; not used in<br>production default |
| Absorption solve | ``block_size`` | 256 | Number of RHS columns<br>processed together; bounds the<br>dense solution buffer, does not<br>change the mathematical<br>system |

Data provenance: ``src/sfate/graph.py``, ``src/sfate/fate.py``, ``inspect.signature(pynndescent.NNDescent.__init__)`` (pynndescent 0.6.0).

**Table S3.** Benchmark-tier metadata

| Tier | n | Cells | Type | Repetitions | Warm-up / cache | Reported metric | Graph build time (s) | RSS peak (MB) | nnz | nnz/(n·k) | Full-pipeline note |
| --- | --- | --- | --- | --- | --- | --- | --- | --- | --- | --- | --- |
| fixture_5k_real | 5,000 | Real microglia (mouse 5xFAD/Cd28-cKO subset) | $\geq 3$ | No separate warm-up; fixed seed 20260909 | Maximum over repetitions | 1.9 | 762.1 | 251,140 | 1.6743 | Full solve measured (validation layer) | |
| synthetic_10k | 10,000 | Synthetic branching topology | $\geq 3$ | No separate warm-up; fixed seed 20260909 | Maximum over repetitions | 3.907 | 859.0 | 480,054 | 1.6002 | Graph-construction tier only | |
| real_75k_full | 74,984 | Full real microglia atlas | $\geq 3$ for graph construction | No separate warm-up; first end-to-end run reported separately | Maximum over repetitions for graph tier; single end-to-end run for full pipeline | 18.444 | 1,413.1 | 3,807,850 | 1.6927 | One first end-to-end run ($\approx 1.76$ GiB peak); reproducible M1 graph benchmark $\geq 3$ reps | |
| synthetic_500k | 500,000 | Synthetic branching topology | $\geq 3$ | No separate warm-up; fixed seed 20260909 | Maximum over repetitions | 66.162 | 4,327.3 | 25,395,928 | 1.6931 | Graph-construction tier only; full-pipeline absorption solve attempted once under default column-wise GMRES and did not converge for the<br>first<br>absorbing<br>cell<br>within<br>49,904<br>iterations<br>(true<br>residual<br>1.09e-5 at<br>rtol = 1e-6) | |

Data provenance: `output/benchmark\_m1\_smoke.json`, `output/benchmark\_m1\_smoke.md`, `EXPERIMENT\_LOG.md`.

## References

1. Anderson, E., et al.. LAPACK Users’ Guide. 1999. doi:10.1137/1.9780898719604 [anderson1999lapack]

2. Pinar Ayata et al.. Lymphoid gene expression supports neuroprotective microglia function. Nature. 2025;648(8092):157–165. doi:10.1038/s41586-025-09662-z [ayata2025]

3. Balay, Satish and Abhyankar, Shrirang and Adams, Mark F., et al.. PETSc/TAO Users Manual. Argonne National Laboratory. 2026(ANL-21/39 - Revision 3.25). doi:10.2172/2998643 [balay2026petsc]

4. Benzi, Michele. Preconditioning Techniques for Large Linear Systems: A Survey. Journal of Computational Physics. 2002;182(2):418–477. doi:10.1006/jcph.2002.7176 [benzi2002survey]

5. Bergen, Volker et al.. Generalizing RNA velocity to transient cell states through dynamical modeling. Nature Biotechnology. 2020;38(12):1408--1414. doi:10.1038/s41587-020-0591-3 [bergen2020scvelo]

6. Berman, Abraham and Plemmons, Robert J.. Nonnegative Matrices in the Mathematical Sciences. . 1994. doi:10.1137/1.9781611971262 [bermanplemmons1994]

7. Brandts, Jan H.. Matlab code for sorting real Schur forms. Numerical Linear Algebra with Applications. 2002;9(3):249--261. doi:10.1002/nla.274 [brandts2002schur]

8. CellRank developers. CellRank installation documentation. . 2026. https://cellrank.readthedocs.io/en/latest/installation.html. doi: [cellrankinstallation]

9. Edmond Chow and Yousef Saad. Experimental Study of ILU Preconditioners for Indefinite Matrices. Journal of Computational and Applied Mathematics. 1997;86(2):387--414. doi:10.1016/s0377-0427(97)00171-4 [chowsaad1997]

10. Deczkowska, Aleksandra et al.. Disease-Associated Microglia: A Universal Immune Sensor of Neurodegeneration. Cell. 2018;173(5):1073–1081. doi:10.1016/j.cell.2018.05.003 [deczkowska2018dam]

11. Deuflhard, Peter and Weber, Marcus. Robust Perron cluster analysis in conformation dynamics. Linear Algebra and its Applications. 2005;398:161–184. doi:10.1016/j.laa.2004.10.026 [deuflhard2005pcca]

12. Dong, Wei and Moses, Charikar and Li, Kai. Efficient k-nearest neighbor graph construction for generic similarity measures. Proceedings of the 20th International Conference on World Wide Web (WWW). 2011:577–586. doi:10.1145/1963405.1963487 [dong2011nndescent]

13. Frank, Anna-Simone and Sikorski, Alexander and Röblitz, Susanna. Spectral clustering of Markov chain transition matrices with complex eigenvalues. Journal of Computational and Applied Mathematics. 2024;444:115791. doi:10.1016/j.cam.2024.115791 [frank2024spectral]

14. Herman, Josip S. and Sagar and Grün, Dominic. FateID infers cell fate bias in multipotent progenitors from single-cell RNA-seq data. Nature Methods. 2018;15(5):379--386. doi:10.1038/nmeth.4662 [herman2018fateid]

15. Hernandez, Vicente and Roman, Jose E. and Vidal, Vicente. SLEPc: A scalable and flexible toolkit for the solution of eigenvalue problems. ACM Transactions on Mathematical Software. 2005;31(3):351--362. doi:10.1145/1089014.1089019 [hernandez2005slepc]

16. Nicholas J. Higham. Accuracy and Stability of Numerical Algorithms. . 2002. doi:10.1137/1.9780898718027 [higham2002accuracy]

17. Keren-Shaul, Hadas et al.. A Unique Microglia Type Associated with Restricting Development of Alzheimer’s Disease. Cell. 2017;169(7):1276–1290.e17. doi:10.1016/j.cell.2017.05.018 [kerenshaul2017dam]

18. Susanne Krasemann et al.. The TREM2-APOE Pathway Drives the Transcriptional Phenotype of Dysfunctional Microglia in Neurodegenerative Diseases. Immunity. 2017;47(3):566--581.e9. doi:10.1016/j.immuni.2017.08.008 [krasemann2017]

19. La Manno, Gioele et al.. RNA velocity of single cells. Nature. 2018;560(7719):494--498. doi:10.1038/s41586-018-0414-6 [lamanno2018velocity]

20. Lange, Marius et al.. CellRank for directed single-cell fate mapping. Nature Methods. 2022;19(2):159--170. doi:10.1038/s41592-021-01346-6 [lange2022cellrank]

21. Lopez, Romain et al.. Deep generative modeling for single-cell transcriptomics. Nature Methods. 2018;15(12):1053--1058. doi:10.1038/s41592-018-0229-2 [lopez2018scvi]

22. Plemmons, Robert J.. M-matrix characterizations. I---Nonsingular M-matrices. Linear Algebra and its Applications. 1977;18(2):175–188. doi:10.1016/0024-3795(77)90073-8 [plemmons1977]

23. Reuter, Bernhard et al.. Generalized Markov State Modeling Method for Nonequilibrium Biomolecular Dynamics. Journal of Chemical Theory and Computation. 2018;14(7):3579--3594. doi:10.1021/acs.jctc.8b00079 [reuter2018gpcca]

24. Reuter, Bernhard and Fackeldey, Konstantin and Weber, Marcus. Generalized Markov modeling of nonreversible molecular kinetics. The Journal of Chemical Physics. 2019;150(17):174103. doi:10.1063/1.5064530 [reuter2019gpcca]

25. Röblitz, Susanna and Weber, Marcus. Fuzzy Spectral Clustering by PCCA+: Application to Markov State Models and Data Classification. Advances in Data Analysis and Classification. 2013;7(2):147--179. doi:10.1007/s11634-013-0134-6 [roblitz2013pcca]

26. Ethan R. Roy et al.. Type I interferon response drives neuroinflammation and synapse loss in Alzheimer disease. Journal of Clinical Investigation. 2020;130(4):1912--1930. doi:10.1172/jci133737 [roy2020ifn]

27. Saad, Youcef and Schultz, Martin H.. GMRES: A generalized minimal residual algorithm for solving nonsymmetric linear systems. SIAM Journal on Scientific and Statistical Computing. 1986;7(3):856--869. doi:10.1137/0907058 [saad1986gmres]

28. Saad, Yousef. ILUT: A dual threshold incomplete LU factorization. Numerical Linear Algebra with Applications. 1994;1(4):387--402. doi:10.1002/nla.1680010405 [saad1994ilut]

29. Setty, Manu et al.. Characterization of cell fate probabilities in single-cell data with Palantir. Nature Biotechnology. 2019;37(4):451--460. doi:10.1038/s41587-019-0068-4 [setty2019palantir]

30. Sharpe, Daniel J. and Wales, David J.. Nearly reducible finite Markov chains: Theory and algorithms. The Journal of Chemical Physics. 2021;155(14):140901. doi:10.1063/5.0060978 [sharpewales2021]

31. V. Simoncini and E. Gallopoulos. Convergence Properties of Block GMRES and Matrix Polynomials. Linear Algebra and its Applications. 1996;247:97--119. doi:10.1016/0024-3795(95)00093-3 [simoncini1996block]

32. Stassen, Shobana V. et al.. Generalized and scalable trajectory inference in single-cell omics data with VIA. Nature Communications. 2021;12(1):5528. doi:10.1038/s41467-021-25773-3 [stassen2021via]

33. G. W. Stewart. A Krylov--Schur Algorithm for Large Eigenproblems. SIAM Journal on Matrix Analysis and Applications. 2002;23(3):601--614. doi:10.1137/s0895479800371529 [stewart2002krylovschur]

34. S. Sukmanyuk and D. Zheltkov and B. Valiakhmetov. Generalized Minimal Residual Method for Systems with Multiple Right-Hand Sides. . 2024. doi:10.48550/arxiv.2408.05513 [sukmanyuk2024]

35. Titley-Peloquin, D. and Pestana, J. and Wathen, A. J.. GMRES convergence bounds that depend on the right-hand-side vector. IMA Journal of Numerical Analysis. 2014;34(2):462--479. doi:10.1093/imanum/drt025 [titleypeloquin2014]

36. Virshup, Isaac et al.. anndata: Access and store annotated data matrices. Journal of Open Source Software. 2024;9(101):4371. doi:10.21105/joss.04371 [virshup2024anndata]

37. Weiler, Philipp et al.. CellRank 2: unified fate mapping in multiview single-cell data. Nature Methods. 2024;21(7):1196--1205. doi:10.1038/s41592-024-02303-9 [weiler2024cellrank2]

38. Wolf, F. Alexander and Angerer, Philipp and Theis, Fabian J.. SCANPY: large-scale single-cell gene expression data analysis. Genome Biology. 2018;19:15. doi:10.1186/s13059-017-1382-0 [wolf2018scanpy]

